# BRD8 guards the pluripotent state by sensing and maintaining histone acetylation

**DOI:** 10.1101/2024.03.20.585857

**Authors:** Li Sun, Xiuling Fu, Zhen Xiao, Gang Ma, Yibin Zhou, Haoqing Hu, Liyang Shi, Dongwei Li, Ralf Jauch, Andrew Paul Hutchins

## Abstract

Epigenetic control of cell fates is a critical determinant to maintain cell type stability and permissive differentiation. However, the epigenetic control mechanisms are not well understood. Here, we show that the histone acetyltransferase reader protein BRD8 impairs the conversion of primed mouse EpiSCs (epiblast stem cells) to naïve mouse ESCs (embryonic stem cells). BRD8 works by maintaining histone acetylation on promoters and transcribed gene bodies. BRD8 is responsible for maintaining open chromatin at somatic genes, and histone acetylation at naïve-specific genes. When *Brd8* expression was reduced, chromatin accessibility was unchanged, but histone acetylation at primed-specific genes was reduced. Conversely, naïve-specific genes had reduced repressive chromatin marks, and acquired accessible chromatin more rapidly during the cell type conversion. We show that this process requires active histone deacetylation to promote the conversion of primed to naïve. Our data supports a model for BRD8 reading histone acetylation to accurately localize the genome-wide binding of the histone acetyltransferase KAT5. Overall, this study shows how the reading of the histone acetylation state by BRD8 maintains cell type stability.

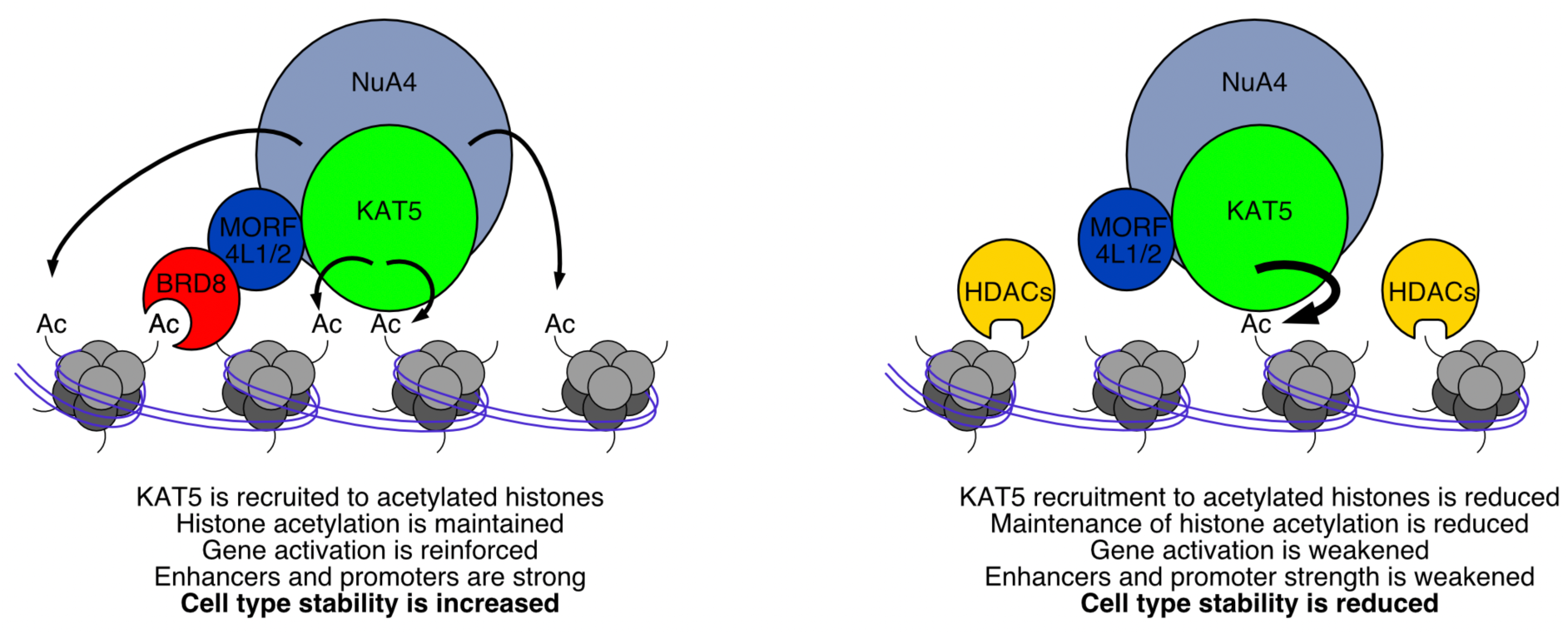

**Key findings:**

- BRD8 blocks the primed-to-naïve transition.
- Reduced *Brd8* promoted the suppression of somatic and primed genes through reduced chromatin repression.
- Reduced *Brd8* enabled the accelerated activation of naïve-specific genes by opening chromatin.
- BRD8 anchors the histone acetyltransferase KAT5 to acetylated genomic loci.

## Introduction

Despite containing the same DNA sequence cells can execute specialized functions and form unique cell types. Changes in cell type are prevalent during embryonic development when the fertilized zygote goes through an elaborate developmental program as cells become progressively restricted to specific cell fates ^1^. Mechanistic control of cell fates remains unclear but is thought to be caused by changes in epigenetic control of DNA and chromatin driven by the activity of cell type-specific transcription factors (TFs) ^2, 3^. However, whilst some TFs are expressed in a cell type-specific pattern, they cooperate with other cell type-independent TFs and epigenetic factors that are often constitutively expressed ^4^. As a consequence, epigenetic factors that erect barriers to cell-type conversions are hard to identify and epigenetic barriers that resist cell-type conversion remain only sporadically described.

Much work has been devoted to the reprogramming of somatic cells to induced pluripotent stem cells, which has revealed a wide array of epigenetic barriers ^5^. However, this somatic cell reprogramming is a major cell type conversion with many changes as it spans essentially nearly all of development. A more amenable model is the interconversion of naïve embryonic stem cells (ESCs) and primed epiblast stem cells (EpiSCs) in mouse. The naïve ESCs resemble the early pre-implantation blastocyst, whilst the primed EpiSCs are more reminiscent of the late epiblast post-implantation and early gastrulation ^6, 7^. Naïve and primed cells have distinct culture media signalling requirements, with naïve cells requiring BMPs and LIF, and primed cells needing Activin A and FGF-signalling ^8^. Attempting to culture naïve cells in primed conditions or *vice versa* results in a mixture of differentiation and/or cell death.

Interconversion of naïve and primed cell types can nonetheless be performed by modulating cell culture conditions and in the absence of transgenes. The process is relatively efficient going from naïve-to-primed, but usually inefficient going from primed-to-naïve ^8^. Naïve and primed states share overlapping gene regulatory modules, for example, OCT4 and SOX2 are active in both cell types ^9–12^. Yet they also have divergent regulatory programs, for example, the TFs OTX2, OCT6, and JUN are active in EpiSCs ^12, 13^, whilst ESRRB, TfFCP2L1, and KLF2, 4, and 5 are active and specific to ESCs ^14–17^. Potentially there are epigenetic roadblocks that stop the EpiSCs from reverting to an earlier developmental stage ^18, 19^. There is also evidence of divergent routes when EpiSCs convert to ESCs ^11, 20, 21^, and multiple intermediate stages between naïve and primed cell types ^7^. Epigenetic barriers erect roadblocks that impair conversion ^18, 22–24^. However, the full nature of these roadblocks remains unclear, and the mechanisms that fix cell fate and resist conversions remain incompletely understood ^2^.

In this study, we set out to understand the epigenetic barriers that prevent the cell fate conversion from primed EpiSCs to naïve ESCs. We focused on the bromodomain family of epigenetic co-factors as several family members have been implicated in cell type conversions ^25–27^. We discovered a role for the bromodomain-containing protein BRD8 in blocking cell-type conversions. BRD8 is a histone acetyltransferase reader protein ^28, 29^, and is a member of the NuA4 protein complex that has histone acetyltransferase activity to promote gene expression ^30^. Reduced *Brd8* promoted the transition of EpiSCs to ESCs in the primed-to-naïve conversion. BRD8 binds to naïve-specific gene loci, modulates epigenetic modifications at those loci, and then influences cell type transition. This effect was associated with changes in the epigenetic state of the cells caused by the loss of the activatory chromatin marks H3K27ac and H3K4me1, and the reduction of the repressive chromatin marks H3K27me3 and H3K9me3 in the transcribed regions of primed-specific genes. Mechanistically this required the action of histone deacetylases and a catalytically active histone acetyltransferase KAT5 (TIP60).

## Results

### *Brd8* impairs the conversion of EpiSCs to ESCs

To explore the epigenetic factors that control the cell fate conversion of EpiSCs to ESCs, we designed a small-scale screen to capture genes that influence the conversion. First, we took advantage of the OG2-GFP EpiSCs. These cells have multiple copies of the *Pou5f1* locus with GFP (green fluorescent protein) with the EpiSC-specific proximal enhancer deleted ^8, 31, 32^. Due to the deletion of the proximal enhancer, they express GFP only when the naïve pluripotency gene expression network is active ^20^. We next modified the culture conditions of the primed-to-naive conversion to reduce the efficiency, so that they are still capable of converting to ESCs, but do so inefficiently to make it easier to identify factors that improve the conversion. We knocked down all bromodomain-family genes that were expressed in ESCs and EpiSCs individually (**Figure 1a, b**). We then converted the cells to the naïve state (**Figure 1c**), and measured the percentage of GFP+ ESCs by FACS (**Figure 1d, and Supplementary** Figure 1a). Note that the knockdown of *Brd4* and *Brd7* was lethal in the primed-to-naïve conversion system and no cells remained after 4 days. This is in agreement with a previous study which showed knockdown of *Brd4* is incompatible with the pluripotent state ^27^. Several bromodomain family factors influenced the reversion of EpiSCs to ESCs, including *Brd8*, *Brd9,* and *Brdt* (**Figure 1d**). The last was surprising as *Brdt* is expressed at low levels in both EpiSCs and ESCs (**Figure 1a**). BRD9 has previously been implicated in controlling human pluripotency through a non-canonical BAF protein complex ^26^, and its inhibition promoted the conversion of ESCs to an EpiSC-like state ^25^. Our data suggests that reduced *Brd9* promotes the conversion of EpiSCs to ESCs. However, a function for *Brd8* in embryogenesis has not been reported in any model system, hence we focused on *Brd8*. Overall, the knockdown of *Brd8* caused the clearest improvement in the generation of GFP+ ESC-like cells (**Figure 1e, f**). This agrees with genome-wide knockout/knockdown studies for the conversion of EpiSCs to ESCs, which placed *Brd8* as either rank 6 ^19^, or rank 678 ^9^ as the best sgRNAs in their screens.

**Figure 1.**
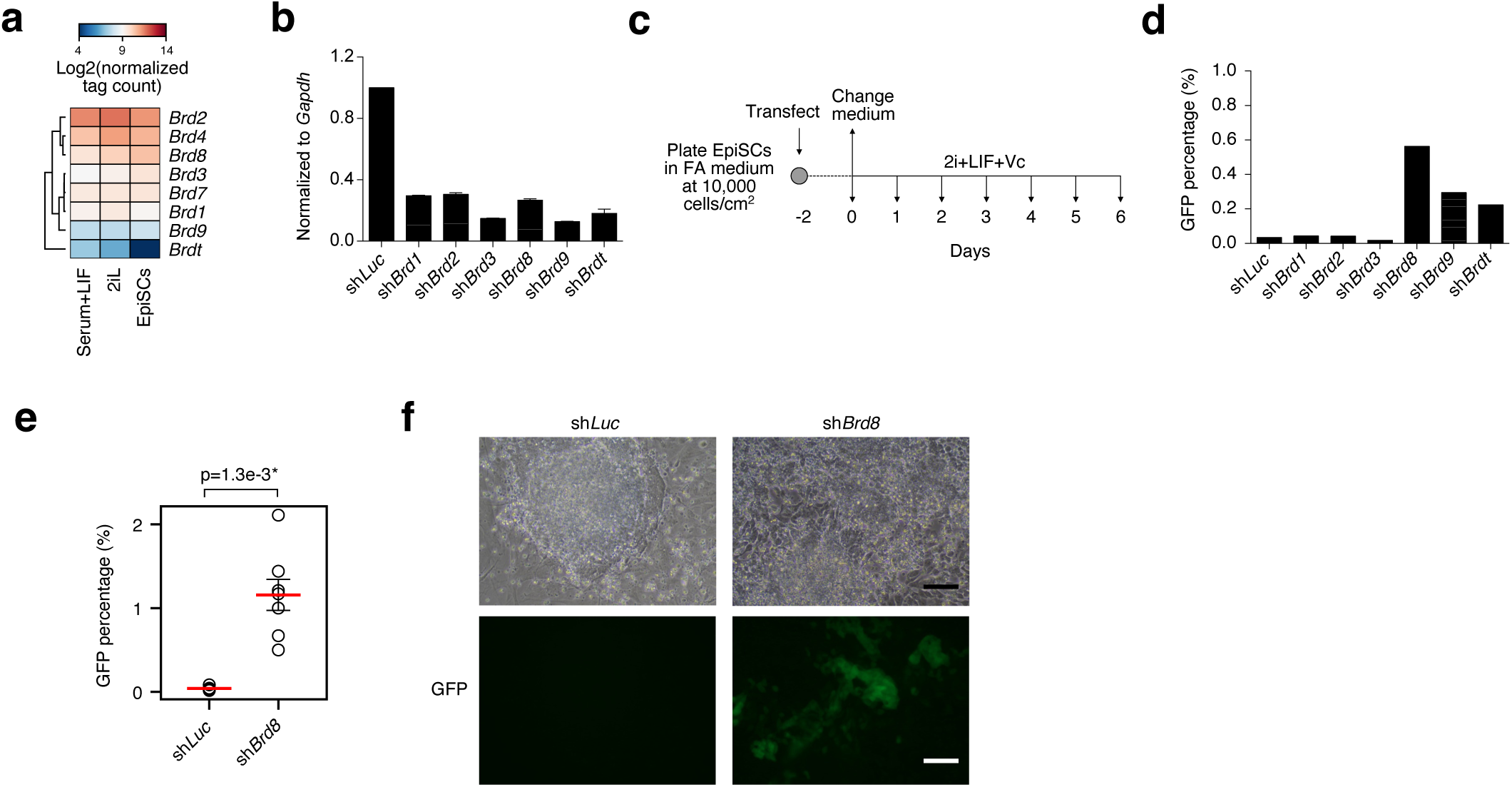
– Bromodomain proteins block the conversion of EpiSCs to ESC. **a.** Heatmap of the expression levels of Bromodomain-family proteins in ESCs grown in Serum+LIF or 2iLIF, and in EpiSCs. **b.** RT-qPCR for the indicated bromodomain-proteins, in the indicated shRNA knockdowns in EpiSCs. Levels of RNA are normalized to *Gapdh*. This experiment was repeated three times. Error bars are standard error of the means. **c.** Schematic of the reprogramming of EpiSCs to ESCs in the primed-to-naïve transition. The EpiSCs used for the conversion contain the OG2-GFP reporter which is only activated in ESC state. **d.** Percent of GFP+ cells as counted by FACS (Fluorescence activated cell sorting) on day 6 of the conversion of EpiSCs to ESCs in the indicated knockdowns. **e.** Dot plot showing the GFP percentages of cells at day 5 of a primed-to-naïve transition in cells transfected with a control sh*Luc*, or an shRNA targeting *Brd8*. The experiment was performed seven times in biological replicate. Significance is from a two-sided Welch’s t-test. * indicates a p-value < 0.05. **f.** Bright field and GFP images of colonies in the indicated knockdowns at day 5 of the primed-to-naïve conversion. Scale bar = 20 μm.

### Loss of *Brd8* promotes the suppression of somatic genes and expression of naïve-specific genes

We next examined in more detail the conversion of EpiSCs to ESCs when *Brd8* was reduced. Time course RNA-seq in cells transfected with sh*Luc* or sh*Brd8* confirmed the up-regulation of naive-specific genes when *Brd8* was knocked down, particularly at days 2-6 (**Figure 2a, b**). Specific marker genes for the naïve state were up-regulated as early as day 4, for example, *Tfcp2l1*, *Dppa5a*, *Dppa2,* and *Esrrb* were all upregulated in the *Brd8* knockdown cells (**Supplementary** Figure 1b and c). Some primed-specific genes were down-regulated slightly earlier in the *Brd8* knockdown cells, for example, *T*, *Dchs1*, and *Otx2* were down-regulated earlier. However, other primed-specific genes, *Ets1* or *Fgf8*, were similar between the two knockdowns. (**Figure 2a, b and Supplementary** Figure 1c and d). Interestingly, principal component analysis (PCA) of the RNA-seq data indicated that whilst the overall trajectory is accelerated in the sh*Brd8* transfected cells, the change is already apparent upon shRNA transfection at day 0, which is 2 days after the addition of the shRNA, but before a change in cell culture medium (**Figure 1b**, **and 2c**). We wondered if this day 0 change was representative of a general suppression of primed-specific genes and was a systematic phenomenon. Hence, we defined sets of primed– and naïve-specific genes by looking at genes that were 2-fold up or downregulated between ESCs and EpiSCs (**Supplementary** Figure 2a**, b and Supplementary Table 1**). The expression of these two gene sets supported an accelerated up-regulation of naïve-specific genes, and accelerated suppression of primed-specific genes (**Figure 2d**). Indeed, primed-specific genes were consistently downregulated at all time points, whilst naive-specific genes were only upregulated at day 6 (**Figure 2d**). This was supported by GSEA, as the down-regulated genes in the sh*Brd8*-transfected day 0 cells were associated with GO terms related to differentiation such as ‘Cell fate commitment’, and neuron, heart, and spinal cord differentiation (**Figure 2e and Supplementary** Figure 2c). Interestingly, GSEA at day 0 for the up-regulated genes suggested blastocyst formation, LIF response, and regulation of stem cell populations as significantly up-regulated (**Supplementary** Figure 2c). This suggests that even as early as day 0, the EpiSCs have suppressed somatic genes, and activated embryonic genes.

**Figure 2.**
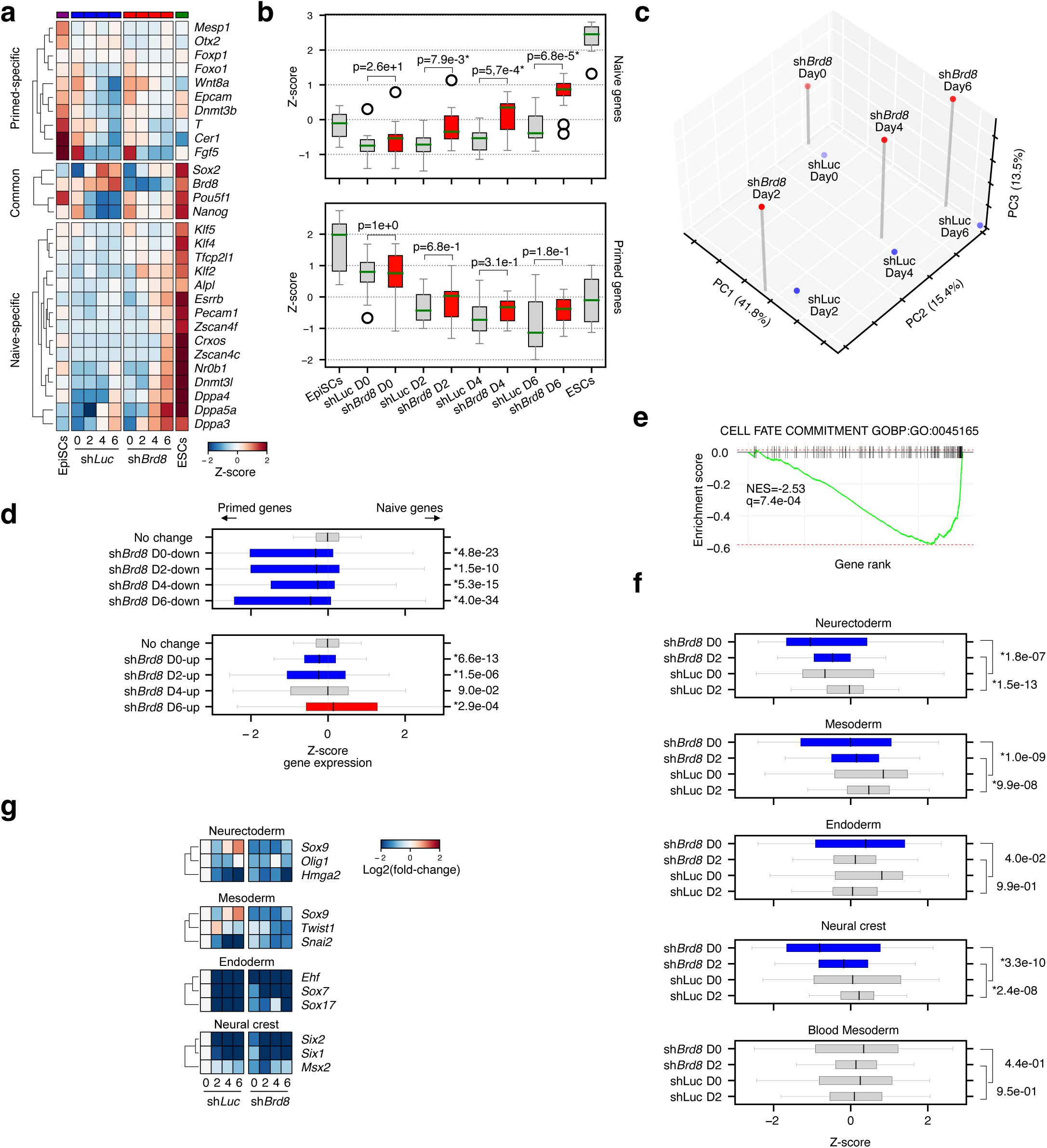
– Knockdown of *Brd8* causes the accelerated downregulation of primed and somatic genes and the late-stage activation of naïve-specific genes. **a.** Heatmap showing RNA-seq data for a selection of marker genes in EpiSCs, 2iL grown ESCs, and in a primed-to-naïve conversion time course in cells transfected with sh*Luc* as a control or sh*Brd8*. **b.** Boxplots of the marker genes’ expression levels, as in **panel a**. Significance is from a two-sided Welch’s t-test. * indicates p<0.01. **c.** Principal component analysis (PCA) of the RNA-seq time course for the primed-to-naïve conversion in sh*Luc* or sh*Brd8*-transfected cells. **d.** Boxplots of the sum of Z-scores for primed or naïve-specific genes (as defined in **Supplementary** Figure 2a and b) versus down-regulated (top boxplot) or up-regulated (bottom boxplot) genes in the sh*Brd8* knockdown. Significance is from a two-sided Welch’s t-test versus the genes that had no change. * indicates a p-value < 0.05. **e.** GSEA enrichment plot for the down-regulated genes at day 0 of the primed-to-naïve conversion in sh*Luc* vs sh*Brd8* cells. A term was considered significant if it had an absolute normalized enrichment score of at least 1.5 and a q-value of <0.01. **f.** Boxplots showing sums of Z-scores for the indicated germ lineage-specific genes as defined in ^4^. Significance is from a two-sided Welch’s t-test versus the genes that had no change. * indicates a p-value < 0.05. **g.** Heatmap of the fold-change for selected germ lineage-specific genes. Fold-change is calculated relative to day 0 sh*Luc* transfected cells.

EpiSCs are poised to differentiate and express low levels of differentiation-related and somatic genes, particularly genes expressed early in gastrulation. To confirm that somatic genes are reduced in the *Brd8* knockdowns we utilized our datasets of germ layer-specific genes ^4^, and scored the overall level of expression of these gene sets in the *Luc* controls and *Brd8* knockdowns (**Figure 2f**). Genes for most germ lineage-specific genes were significantly downregulated in the sh*Brd8* knockdowns at early time points (**Figure 2f**). These results are exemplified by the downregulation of germ layer-specific genes as early as day 0 in the *Brd8* knockdown cells (**Figure 2g**). Overall, this data suggests that reduced *Brd8* destabilizes the expression of primed and differentiation-specific genes in the earliest stages of conversion to ESCs which is succeeded by an acceleration in the activation of naïve-specific genes in the late stage. We conclude that BRD8 acts as an epigenetic rheostat to maintain cell fates whilst its depletion derails the status quo.

### Reduced *Brd8* promotes the opening of chromatin at naïve-specific genes

To explore the changes in the chromatin state, which can often precede changes in gene expression ^33^, we assayed chromatin accessibility using ATAC-seq ^34^, and defined open and closed groups as previously described ^33, 35^. The resulting pattern of chromatin changes was complex (**Figure 3a and Supplementary** Figure 3a). We focused on those peaks that were dynamically changing over the time course, which amounted to 36,203 loci. We divided those peaks into several categories based on their pattern in the sh*Brd8* knockdown (**Figure 3b and c**). Knocking down *Brd8* had varied effects on chromatin accessibility and the changes were equally distributed between accelerated and decelerated opening or closing (**Figure 3c**). This was curious and suggests that reduced *Brd8* is increasing overall cellular plasticity, rather than influencing the chromatin for a specific cell type. To explore this further, we associated the nearby genes with their expression level in naïve or primed cells using a Z-score (**Figure 3b and c**). This will reveal if a category of chromatin changes is associated with naïve or primed-specific genes. Despite the major changes in chromatin, only one category of chromatin changes was significantly associated with naïve-specific genes, sh*Brd8*-specific open loci (**Figure 3b and c**). This is reflected in the accessibility of chromatin at naïve-specific genes, such as *Zfp42*, *Dppa5a, App4, and Nr0b1* loci that were all open at day 6 when *Brd8* was knocked down (**Figure 3d, e and Supplementary** Figure 3b and c). Curiously, no category of chromatin change was significantly associated with primed-specific genes (**Figure 3c**), suggesting that chromatin accessibility is not a driving force in the suppression of somatic genes. Indeed, primed-specific genes tended to retain open chromatin in the *Brd8* knockdowns, and in some cases even had increased chromatin accessibility in the *Brd8* knockdown cells at day 6. For example, the primed genes *Gbp2*, *Dab1*, *Aplpr,* and *Flt1* all had chromatin accessibility that was similar to the control sh*Luc* cells or increased in the *Brd8* knockdowns (**Figure 3d, e, and Supplementary** Figure 3d and e). These results, combined with the RNA-seq results, suggest that chromatin accessibility changes only underly the increases in naïve-specific gene expression. Primed-specific genes, conversely, retain open chromatin when *Brd8* was knocked down. This suggests two distinct epigenetic mechanisms are regulating naïve and primed genes.

**Figure 3.**
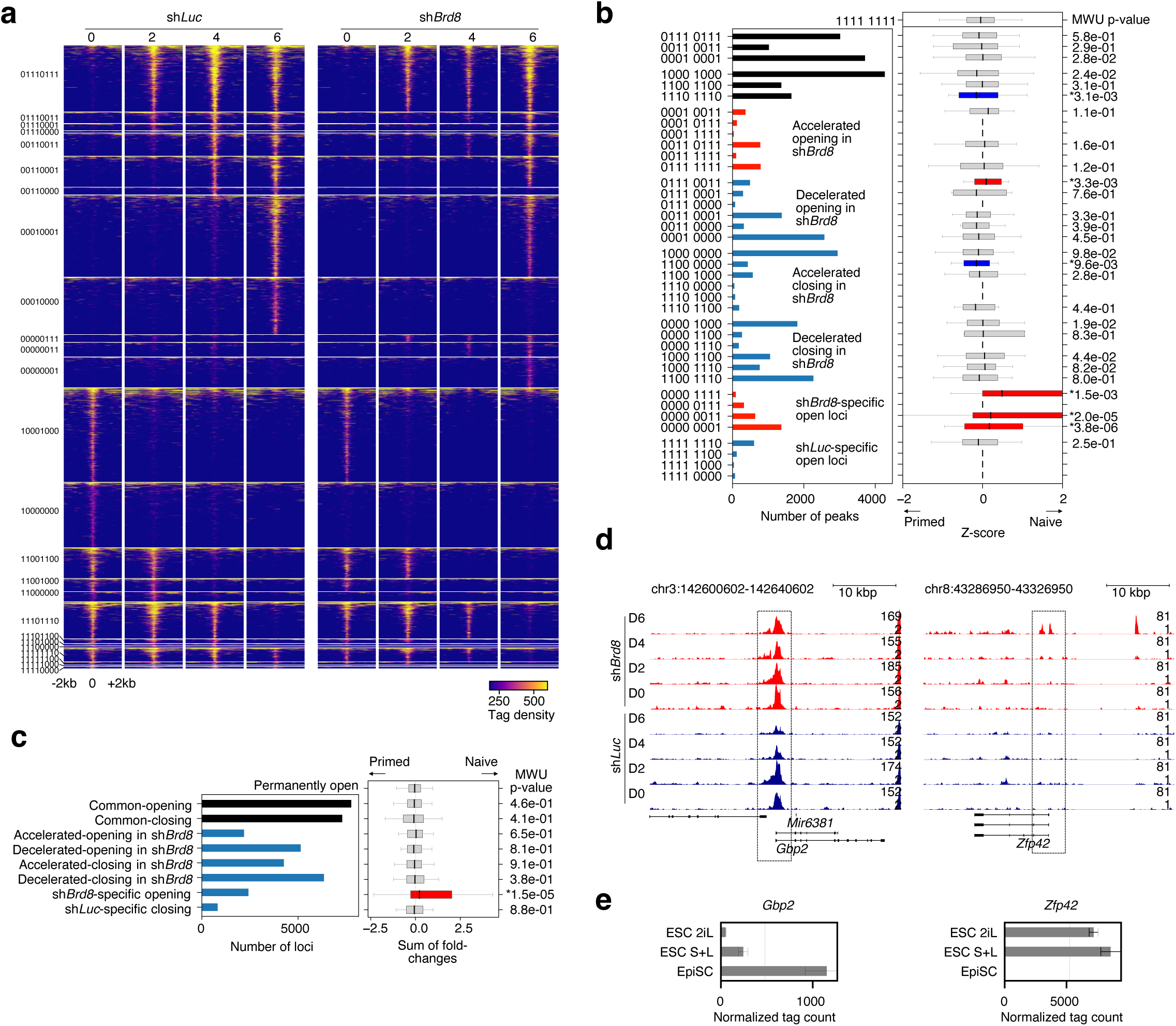
– Reduced *Brd8* expression leads to changes in chromatin accessibility. **a.** Pileup heatmaps for selected ATAC-seq accessibility data showing the clusters of chromatin loci that change in the indicated conditions according to the binary key on the left-hand side. For the binary key, 1 indicates the presence of an open ‘peak’ of binding, whilst 0 indicates no peak detected. The full heatmap showing all clusters is in **Supplementary** Figure 3a. **b.** Combined plot showing the number of peaks in each category of binding (As in **panel a**) (left side), and the expression of genes with a TSS within 2000 bp of the chromatin accessibility locus (right side). The genes were defined as a Z-score based on their expression in EpiSCs versus ESCs (See **Supplementary** Figure 2a and b). A positive Z-score indicates a gene more likely to be high in naïve ESCs, and a negative Z-score more likely to be high in EpiSCs. Significance is from a Mann-Whitney U test. * indicates a p-value < 0.05. **c.** Bar chart showing the number of peaks in the categories defined in **panel b**, and the sum of fold-changes for all genes for naïve versus primed cells. Significance is from a Mann-Whitney U test. * indicates a p-value < 0.05. **d.** Genome pileup plots at the primed gene *Gbp2*, and the naïve gene *Zfp42*. **e.** Bar chart showing gene expression levels of the primed-specific gene *Gbp2* and the naïve-specific gene *Zfp42*, from RNA-seq data for primed and naïve cells grown in 2iLIF (2iL), serum+LIF (S+L) or EpiSC culture conditions.

### BRD8 and the NuA4 complex associates with naïve-specific transcription factors in ESCs

We next looked at the genome-wide binding of BRD8 using CUT&Tag ^36^. Surprisingly, genome-wide BRD8 binding in EpiSCs was a subset of the binding pattern seen in ESCs, with 5,485 peaks common to EpiSCs and ESCs, 5,684 loci specific to ESCs and only 454 loci specific to EpiSCs (**Figure 4a**). Motif discovery in the common and ESC-specific peaks indicated that BRD8 was primarily associated with motifs related to naïve-specific transcription factors, such as TFCP2L1, ESRRB, and PRDM15 ^15, 37,38^ (**Figure 4b**). We reanalyzed ChIP-seq data in ESCs matching the predicted motifs in **Figure 4b**, including the key pluripotent TFs OCT4, SOX2, and KLF4, along with the naïve-specific TFs PRDM15, TFCP2L1 and ESRRB factors in ESCs ^38, 39^. BRD8 was indeed bound at the same loci as TFCP2L1, ESRRB, and PRDM15 in ESCs (**Figure 4c and Supplementary** Figure 4a**, b**). Interestingly, transcription factors that are common to both ESCs and EpiSCs ^10^, OCT4 and SOX2, were not bound at BRD8-bound loci in ESCs or EpiSCs (**Supplementary** Figure 4a**, b**). This suggests that BRD8 specifically binds to loci only with naïve-specific TFs in ESCs. Indeed, reanalysis of ChIP-seq data in EpiSCs showed that BRD8 is not associated with OCT4, OCT6, SOX2, OTX2, and is only bound together with the primed pluripotency marker ZIC2-bound loci (**Figure 4d and Supplementary** Figure 4b**, c**).

**Figure 4.**
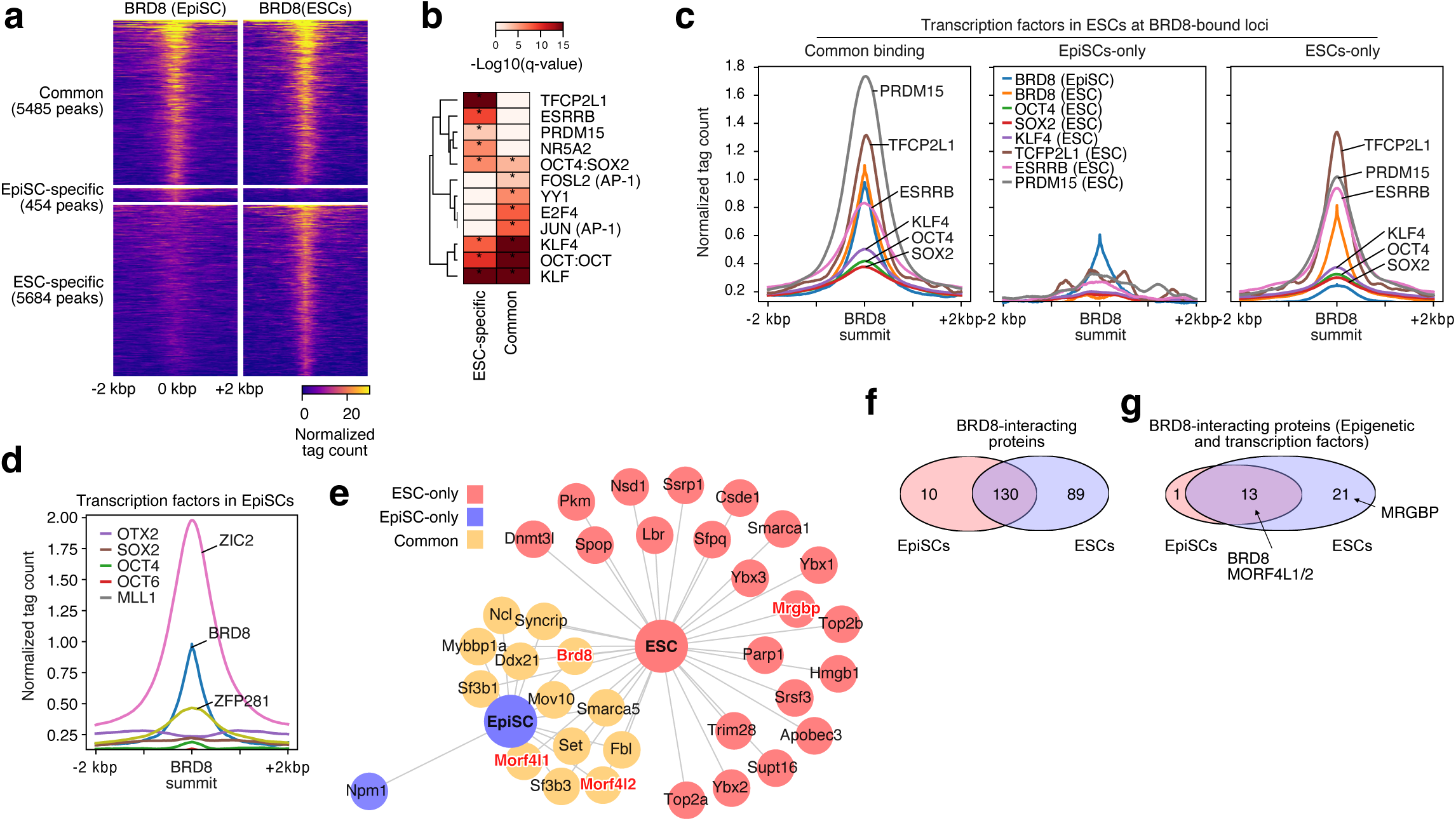
– BRD8 is a member of a novel protein complex in EpiSCs and ESCs, and associates with naïve TFs. **a.** Heatmap pileups for CUT&Tag of BRD8 in EpiSCs and 2iL grown ESCs. Peaks were divided into those loci common to ESCs and EpiSCs (Common-binding), and ESC– and EpiSC-specific loci. **b.** TF motif discovery in the EpiSC-specific group versus the BRD8 common binding peaks. **c.** Cumulative pileup plots for a selection of ESC ChIP-seq data. ChIP-seq data is from GSE11431 ^39^, and GSE73692 ^38^. The data is centered on the BRD8 binding peak for the groups of binding as defined in **panel a**. See also **Supplementary** Figure 4a for the full heatmap of all binding factors. **d.** Cumulative pileup plots for a selection of ESC ChIP-seq data. Data is from GSE74636 ^10^, GSE73992 ^52^, and GSE93042 ^54^. See also **Supplementary** Figure 4b for the full heatmap of all binding factors. **e.** Network of interacting transcription or epigenetic factors detected in the Co-IP/MS for BRD8 binding partners in EpiSCs and ESCs. Commonly detected peptides are marked in orange, red is specific to ESCs, and blue is specific to EpiSCs. The NuA4 complex members Morf4l1/2 and Mrgbp are indicated. The full table of interacting proteins is in **Supplementary** Figure 5a and **Supplementary Table 2**. **f.** Venn diagram of the overlap of all BRD8 interacting proteins in ESCs and EpiSCs. **g.** Venn diagram, as in **panel f**, but only including transcription and epigenetic factors.

To identify the physical interactions of BRD8, we next performed co-immunoprecipitation and mass spectrometry (Co-IP/MS) to identify protein-binding partners for BRD8 in ESCs and EpiSCs. In total we identified 229 significant protein interactions in the two cell types (**Figure 4e-g Supplementary** Figure 5a**, and Supplementary Table 2**). The interacting partners were similar in the two cell types with 130 proteins in common and tended to be related to DNA binding (**Figure 4f and Supplementary** Figure 5b). Interestingly, no naïve-specific TFs were identified in the ESC BRD8-Co-IP/MS data, nor was ZIC2 identified, suggesting that BRD8 is interacting indirectly with these factors (**Figure 4d-f, Supplementary** Figure 5a **and Supplementary Table 2**). BRD8 is a member of the histone acetyltransferase NuA4 complex ^40^, however, only a few NuA4 complex members were identified interacting with BRD8: MORF4L1/2, and MRGBP, the latter of which was only co-precipitated in ESCs (**Figure 4e, g**). These data suggest that BRD8 is a peripheral component of the NuA4 complex and binds MORF4L1/2 in ESCs and EpiSCs, along with MRGBP in ESCs.

### BRD8 binds at promoters and its loss reduces signals associated with active genes

BRD8 binds acetylated histones through its bromodomain and is a component of the NuA4 chromatin-modifying complex ^30, 41, 42^. Hence we expect reduced *Brd8* expression will lead to changes in chromatin. Surprisingly, the western blot of whole cell levels of histone marks did not indicate any changes (**Figure 5a**), suggesting that modulation of chromatin is context-specific and not genome-wide. ATAC-seq accessibility was unchanged at BRD8-bound loci (**Figure 5b**). This was not surprising as we have already shown that loci accessible in primed EpiSCs remain open in the *Brd8* knockdowns (**Figure 3d**). These results suggest that BRD8 mainly impacts gene expression in a mechanism independent of changes in chromatin accessibility.

**Figure 5.**
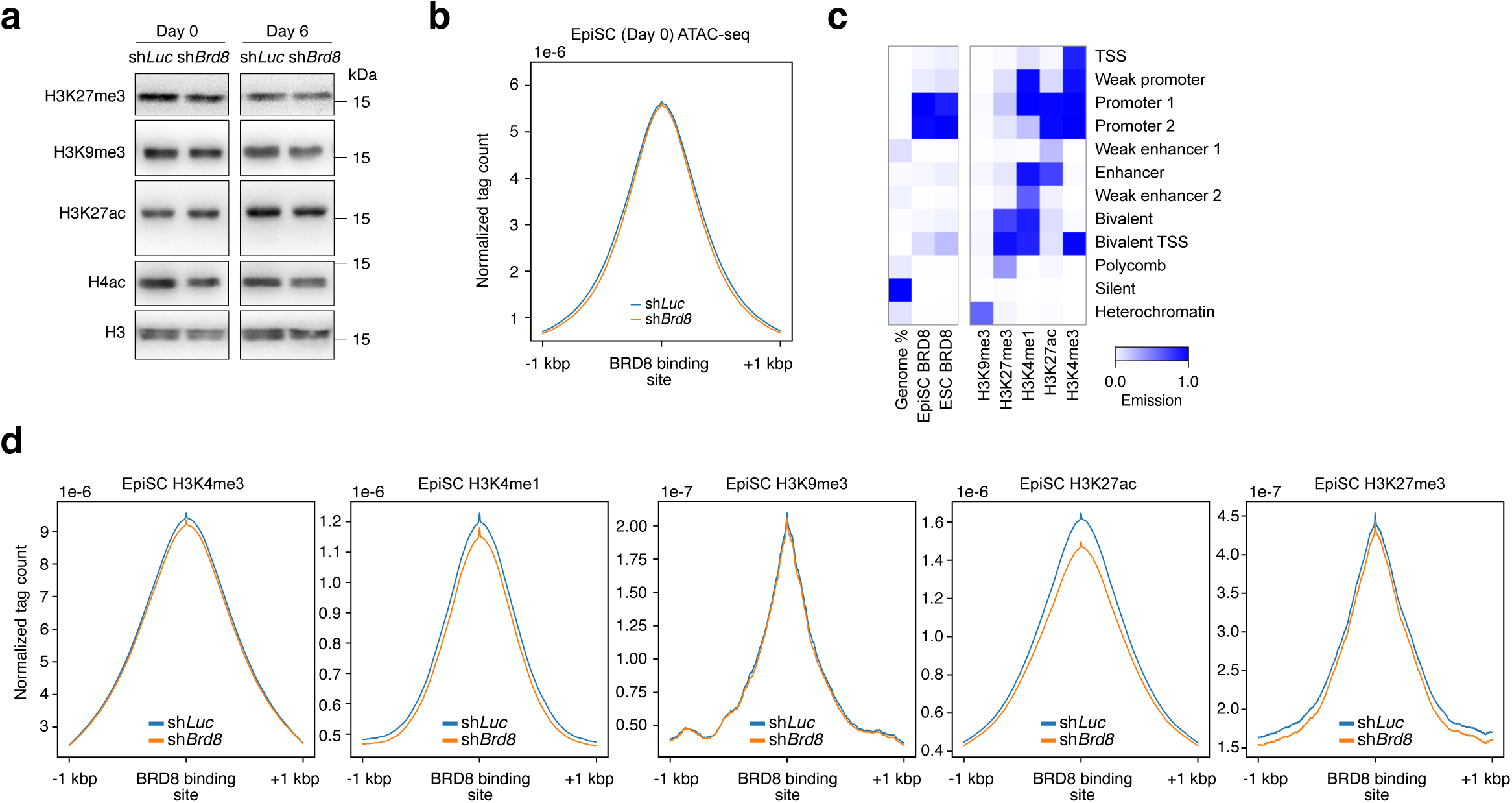
– BRD8 regulates chromatin states in ESCs. **a.** Western blot of histone modifications in day 0 and day 6 EpiSCs transfected with the indicated shRNA, and undergoing a primed-to-naïve transition. **b.** Density pileup for ATAC-seq data centered on BRD8 binding in EpiSCs. ATAC-seq data is from the day 0 data from cells transfected with shRNAs targeting *Luc* (Control) or *Brd8*. **c.** ChromHMM heatmap showing the relative abundance of the factor (left heatmap). The 12-state model (right heatmap) was generated using chromatin ChIP-seq data in the subsequent section. **d.** Density pileup for H3K4me3, H3K4me1, H3K9me3, H3K27ac, and H3K27me3 in EpiSCs transfected with the indicated shRNA. The pileups are centered on the BRD8 binding site in EpiSCs.

To explore this at the histone modification level, we generated genome-wide data for histone modifications that mark promoters (H3K4me3), active promoters (H3K27ac), promoters and enhancers (H3K4me1), repressed polycomb (H3K27me3) and heterochromatin (H3K9me3) in EpiSCs transfected with a control sh*Luc*, or an shRNA targeting *Brd8*. EpiSCs are a relatively underexplored cell type, and we have generated the first genome-wide chromatin map. Hence, we used our histone data to generate an EpiSC-specific ChromHMM 12-state model ^43^, to identify the chromatin features BRD8 is bound to. The ChromHMM model displayed chromatin features that have been observed in other cell types, including transcription start sites (TSSs) (H3K4me3), promoters (H3K4me3, H3K27ac, H3K4me1), enhancers (H3K4me1, H3K27ac), bivalent domains (H3K27me3, H3K4me1), polycomb (H3K27me3) and heterochromatin (H3K9me3) (**Figure 5c**). Using this 12-state model, BRD8 was primarily associated with promoters, and weakly with bivalent promoters (**Figure 5c**).

BRD8 was not bound to heterochromatin regions (**Figure 5c**), and it was unsurprising that H3K9me3 was unchanged upon *Brd8* KD (**Figure 5d**). Similarly, when *Brd8* was knocked down the promoter marker H3K4me3 was not substantially reduced (**Figure 5d**). Instead, the main changes in chromatin marks were in H3K4me1, H3K27ac, and to a lesser extent, H3K27me3 (**Figure 5d**). Interestingly, the decline in H3K7me3 was in the flanking regions, rather than the actual location of BRD8 binding. BRD8 was previously identified in a NuA4-super complex containing PRC2 proteins in sarcoma ^44^, however, we did not detect any PRC2 components in the Co-IP/MS, suggesting changes in H3K27me3 are indirect. As the biggest declines upon *Brd8* depletion were in regions with H3K4me1 and H3K27ac marks (**Figure 5d**), we reasoned that promoters were being decommissioned upon BRD8 removal.

### BRD8 erects a barrier by modulating chromatin modifications inside gene bodies

When *Brd8* expression was reduced, H3K27ac declined, along with H3K4me1 and flanking regions of H3K27me3 (**Figure 5d**). BRD8 tended to concentrate around promoters (**Figure 5c, 6a**), and binding signals were most prominent close to the TSS (**Figure 6a**) but were also enriched in the 5’UTRs and exons of transcribed genes (**Figure 6a**). Pileups of BRD8 across all expressed genes in EpiSCs showed that BRD8 was also present in the transcribed gene bodies (**Figure 6b**). To understand the impact of BRD8 binding on transcription we divided all genes up into expression quartiles, ranked by their total expression level. Based on this, genes were divided into one of five classes: Either not expressed/detectable, or their rank in the expression quartiles (Q1-Q4). We then measured BRD8 binding density across these transcripts.

**Figure 6.**
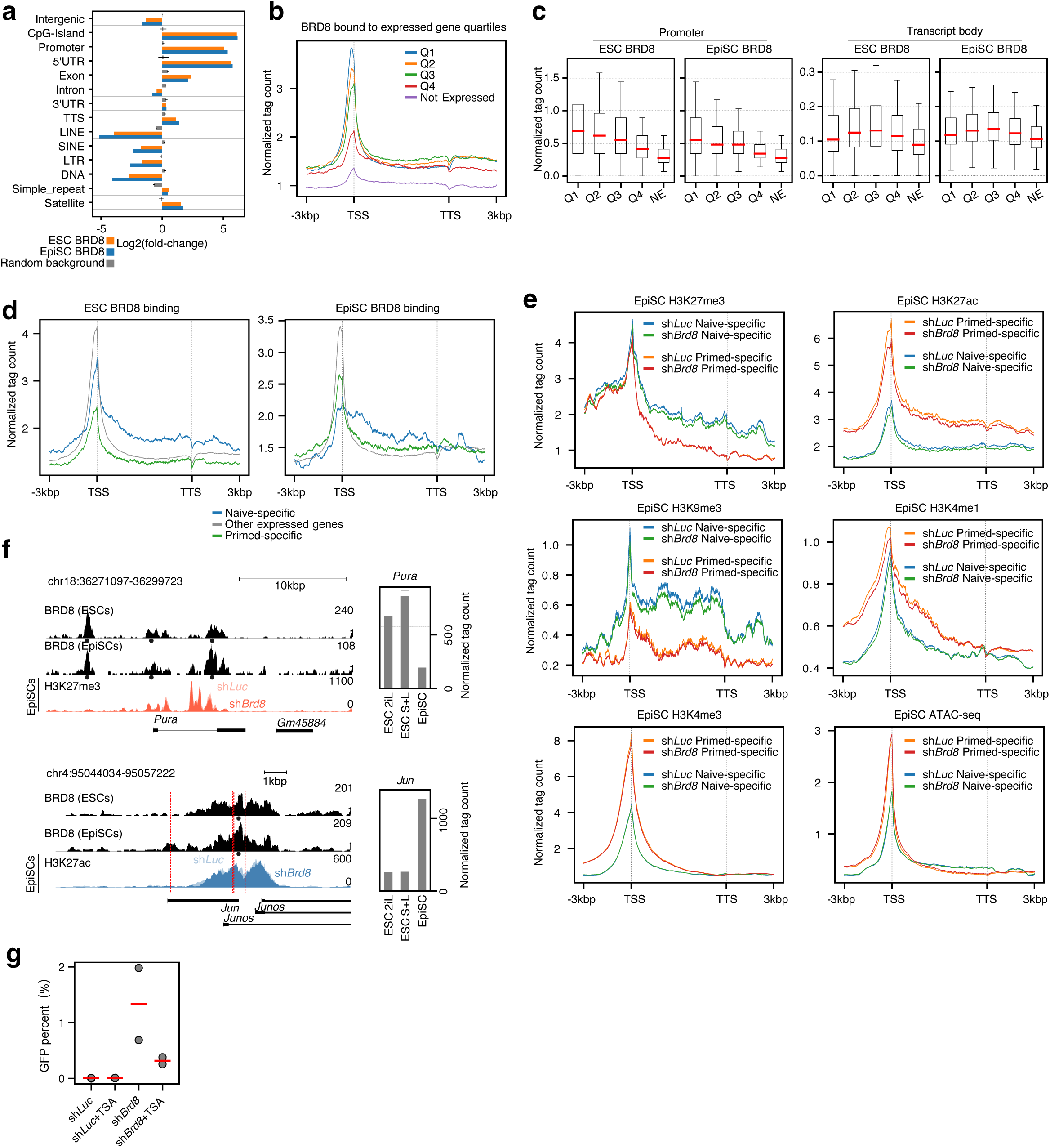
– BRD8 regulates chromatin at naïve and primed-specific genes. **a.** Bar chart of over or under-representation of genome features for the binding of BRD8 in ESCs and EpiSCs. Enrichment was calculated with HOMER ^57^. **b.** A pileup of BRD8 CUT&Tag signals across all genes in the mouse genome. Genes were divided into expressed (normalized tag count >100) and not expressed. The expressed genes were further subdivided into their expression quartiles, Q1-Q4. The windows of binding are centered on the TSS and TTS (transcription termination site) and all transcripts are scaled to a uniform length. The flanking 5’ and 3’ regions 3 kbp from the TSS or TTS are shown. For this and all subsequent pileup plots in this figure. **c.** Box plot showing the ratio of BRD8 signal at the TSS (+/− 500 bp around the TSS) versus the level in the gene body (500 bp 3’ of the TSS and 500 bp 5’ of the TTS). **d.** Pileup of BRD8 CUT&Tag signal across naïve, primed-specific genes versus all other expressed genes (naïve and primed-specific genes are defined in **Supplementary** Figure 2a and b) in ESCs (left plot) or EpiSCs (right plot). **e.** Pileups of H3K4me1, H3K4me3, H3K27ac, H3K9me3, H3K27me3, and ATAC-seq accessibility at naïve and primed-specific genes (naïve and primed-specific genes are defined in **Supplementary** Figure 2a and b). **f.** Genome view showing BRD8 binding at the naïve-specific gene Klf2 (top plot) and the primed gene Jun (bottom plot) in EpiSCs. Gene expression in primed (EpiSCs) and naïve (2iL and SL) ESCs are shown on the right of the genome view. **g.** Effect of the HDAC inhibitor TSA on the conversion of primed EpiSCs to naïve ESCs in a primed-to-naïve conversion. GFP+ cells were counted by FACS on day 6. The experiment was performed in biological duplicate.

At the TSS there was a clear correlation between BRD8 binding and higher levels of expression (**Figure 6b**). However, BRD8 binding in the transcript body was more likely to be associated with moderately expressed genes in the Q2 or Q3 quartiles (**Figure 6b and c**). Even Q4 genes had appreciable levels of BRD8-binding to their transcript bodies, and only silent genes lacked BRD8 binding (**Figure 6b and c**).

Cell type-specifically expressed genes tend to be in the moderate range of overall gene expression levels ^4^, suggesting BRD8 is regulating cell-type-specific genes. Hence we divided genes in ESCs and EpiSCs into naïve or primed-specific and all other genes (**Supplementary** Figure 2a and b) and then measured BRD8 binding at these subsets of genes (**Figure 6d**). The pattern at the TSS and transcribed body was complex: In ESCs, BRD8 binding was high at both the TSS and in the transcript body, whilst in EpiSCs BRD8 binding at the TSS was high, but both naïve and primed-specific genes were bound by BRD8 inside the transcript body (**Figure 6d**). This pattern could be observed at specific genome loci. For example, for the two naïve-specific genes *Jun*, *Nr0b1* and *Tfcp2l1*, BRD8 binding at the TSS was reduced in EpiSCs compared to ESCs, but the BRD8 signal in the transcribed bodies was comparable between the two cell types (**Figure 6f and Supplementary** Figure 6a). Conversely, at two primed-specific genes, *Zic2* and *Pdlim7*, BRD8 binding in ESCs was lower than in EpiSCs in both the TSS and in the transcribed gene body (**Supplementary** Figure 6b).

Considering that BRD8 was also spread across transcribed bodies, we looked at the changes in chromatin across the same naïve and primed gene sets when *Brd8* was knocked down. As in the case for the BRD8-bound loci (**Figure 6d**), activatory marks H3K27ac, and H3K4me1 levels declined across primed-specific transcripts when *Brd8* was knocked down (**Figure 6e**). However, ATAC-seq accessibility and H3K4me3 were unaltered (**Figure 6e**). Intriguingly, repressive marks H3K27me3 and H3K9me3 were unaltered at the TSSs, but were reduced at naïve-specific transcript bodies, for example in the naïve-specific gene *Pura* (**Figure 6f**). These data suggest that the ultimate impact of reduced *Brd8* is to disrupt activatory chromatin marks at primed-specific genes and repressive marks at naïve-specific genes, thus loosening the overall epigenetic stability by weakening chromatin regulation.

These data suggest that the weakening of H3K27ac at primed and naïve genes underlies the ability of reduced *Brd8* to promote the primed-to-naïve transition. Consequently, we reasoned that inhibition of histone deacetylases (HDACs) using the inhibitor TSA would block the action of reduced *Brd8* expression. This was indeed the case, as TSA reverted the rate of GFP+ cells nearly back to the sh*Luc* control levels. (**Figure 6g**). These data indicate that BRD8 is binding to acetylated histones, and when Brd8 expression is reduced active histone deacetylation is required for the primed-to-naïve conversion.

### BRD8 co-operates with the acetyltransferase KAT5 on primed-specific genes

We sought to unravel a mechanism to explain how BRD8 drives the reduction of histone acetylation at primed-specific genes. In our Co-IP/MS data, BRD8 associated with components of the NuA4 complex in EpiSCs (**Figure 4e**). In our MS data, we could not readily identify the acetyltransferase responsible. We surmised that BRD8 might collaborate with KAT5 (TIP60), which is the main catalytic subunit of the NuA4 complex ^30^. We employed a Co-IP Western blot and a Kozak sequence-driven over-expression system to test whether BRD8 and KAT5 could interact (**Figure 7a**). In this system BRD8 could co-precipitate KAT5 (**Figure 7b**), suggesting that KAT5 may be the histone acetyltransferase critical for BRD8 function.

**Figure 7.**
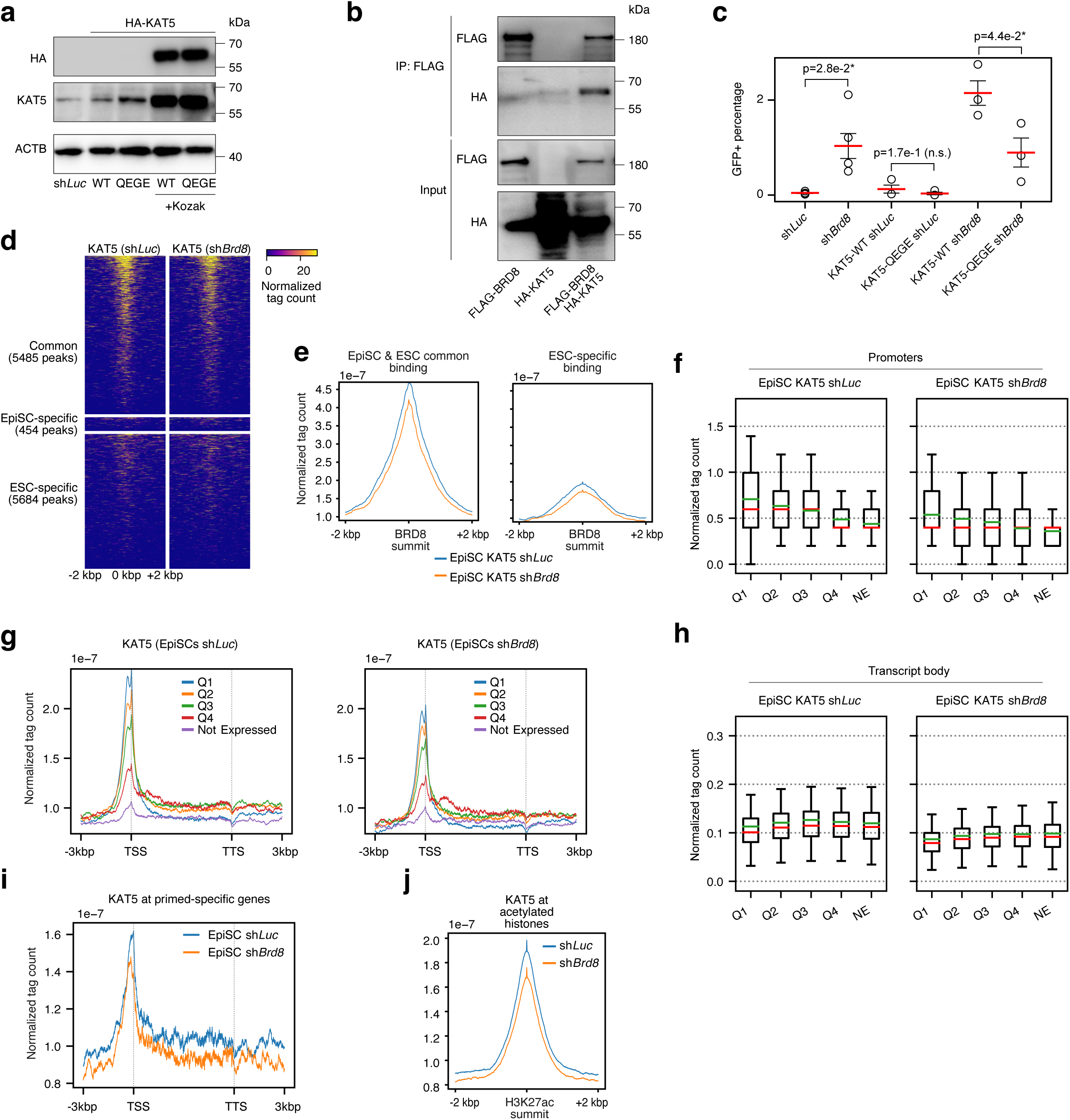
– BRD8 requires KAT5 for activity. **a.** Western blot for the overexpression of KAT5 wildtype (WT) or a catalytic-null KAT5^QEGE^ (KAT5^Q377E/G380E^) 45. **b.** Western blot of Co-IP with the indicated antibodies in cells transfected with HA-tagged KAT5, FLAG-tagged BRD8, or with both vectors. **c.** GFP+ cell counts on day 6 as counted by FACS in cells transfected with the indicated shRNA or with a Kat5 overexpression vector. GFP+ cells counted by FACS at day 6 of a primed-to-naïve-transition. This experiment was performed in biological triplicate. The red bar indicates the mean and the error bars are the standard error of the mean. Significance is from a two-sided Welch’s t-test, * indicates less than 0.05. **d.** Heatmap pileups for CUT&Tag of KAT5 in EpiSCs transfected with a control sh*Luc* or sh*Brd8*. Peaks were divided as in Figure 4A by BRD8 binding. The heatmap is centered on BRD8 binding and extends 2 kbp either side of the peak summit. **e.** Density pileups for KAT5 at BRD8 binding sites in EpiSCs and ESCS (as defined in Figure 4a). The pileups are centered on the BRD8 binding summit. **f.** A pileup of KAT5 CUT&Tag signals across all genes in the mouse genome. Genes were divided into expressed (normalized tag count >100) and not expressed. The expressed genes were further subdivided into their expression quartiles, Q1-Q4. The windows of binding are centered on the TSS and TTS (transcription termination site) and all transcripts are scaled to a uniform length. The flanking 5’ and 3’ regions 3 kbp from the TSS or TTS are shown. **g.** As in **panel f**, but for the transcript bodies, defined as 500 bp 3’ of the TSS and 500 bp 5’ of the transcription end site (TES). **h.** Box plots showing the ratio of KAT5 ChIP-seq read density in the indicated EpiSCs transfected with an shRNA targeting *Luc* or *Kat5*. Boxplots are divided by the reads at the promoter (+/− 500 bp around the TSS) versus the level in the gene body (500 bp 3’ of the TSS and 500 bp 5’ of the TTS). The green line is the mean, the red middle line is the median. **i.** A pileup of KAT5 C&T signals across primed-specific genes (As defined in **Supplementary** Figure 2a and b), in cells transfected with an shRNA targeting *Luc* or *Brd8*. **j.** Pileup of KAT5 binding in cells transfected with a shRNA targeting *Luc* or *Brd8* at all acetylated histones in EpiSCs.

We speculated that BRD8 is responsible for correctly localizing KAT5 to primed-specific genes, and reduced *Brd8* would lead to incorrect localization and a consequent reduction in histone acetylation. Hence, if KAT5 were required for the effect of reduced *Brd8* on the primed-to-naïve conversion then overexpression of *Kat5* would be expected to synergize with a *Brd8* knockdown. This was indeed the case. When *Kat5* was overexpressed in a primed-to-naïve conversion it had only a modest effect in improving the percentage of resulting GFP+ naïve cells (**Figure 7c**). However, the combination of *Kat5* overexpression and *Brd8* knockdown substantially improved the conversion of primed EpiSCs to naïve ESCs (**Figure 7c**). Importantly, this effect required a catalytically active KAT5 as a double mutant catalytically inactive KAT5^Q377E/G380E^ (Kat5^QEGE^) ^45^, was incapable of synergizing with the *Brd8* knockdown (**Figure 7c**). Interestingly, this effect is converse to KAT5’s role in ESCs, where it promotes self-renewal and represses differentiation genes, by a mechanism that does not require lysine acetylation activity ^46^. These results indicate that functional histone acetyltransferase activity is required for BRD8s function.

We next explored the genome binding of KAT5. CUT&Tag of KAT5 in EpiSCs showed that KAT5 was primarily associated with BRD8-bound loci in EpiSCs (**Figure 7d and e**). Importantly, knockdown of *Brd8* reduced the binding of KAT5, indicating that BRD8 is indeed responsible for anchoring KAT5 to the genome (**Figure 7d and e**). We next looked at the association of KAT5 binding with gene expression levels. KAT5 binding tended to be higher at the TSSs of expressed genes in Q1 (**Figure 7f, g**). However, KAT5 binding in the transcript body tended to be higher in Q2-Q4 genes (**Figure g and h**). Knockdown of *Brd8* reduced KAT5 binding at all gene expression quartiles in the promoter (**Figure 7f-h**). Based on these data we proposed that KAT5 maintains chromatin at primed-specific genes. When we measured the level of KAT5 at primed-specific genes it was indeed high (**Figure 7i**). Importantly, when *Brd8* was knocked down KAT5 binding was also reduced at primed-specific genes (**Figure 7j**). Indicating that BRD8 is anchoring KAT5 to both the TSS and transcribed body of primed-specific genes.

Finally, we measured KAT5 binding at acetylated histones, and it was reduced when *Brd8* was knocked down (**Figure 7j**). Overall, these data indicate that BRD8 is responsible for anchoring KAT5 at primed-specific genes and acetylated histones. Reduced *Brd8* expression leads to disruption of the maintenance of acetylation at primed-specific genes, which ultimately leads to increased cell plasticity due to the erosion of epigenetic marks.

## Discussion

Our data suggests that BRD8 has dual roles in primed EpiSCs. Firstly, it maintains acetylation at primed-specific genes, and so stabilizing cell type. Secondly, it primes the chromatin accessibility of naive-specific genes, making them ready for activation. This is accompanied by reduced repressive marks at naïve-specific genes. As a consequence, when *Brd8* expression is reduced EpiSCs become more permissive to the primed-to-naïve conversion.

BRD8 is a member of the NuA4 epigenetic complex, but surprisingly little is known about BRD8’s role. Mechanistically, BRD8 binds to acetylated lysine on histones through a conserved structural motif ^29^, and the best characterized biological roles for BRD8 are in cancer ^47–49^. For example, BRD8 has a key role in maintaining *TP53* (p53) expression in glioblastoma ^42^. Interestingly it represses p53-target genes by blocking p53 binding to the genome through maintaining compact chromatin. This is opposite to the pattern seen here, where reduced *Brd8* leads to reduced histone acetylation and reduced histone methylation, although the latter appears indirect. BRD8 also has a role in regulating cell cycle progression in colorectal cancer ^50^.

Intriguingly BRD8 performs this function independently of the wider NuA4 protein complex. Hence, the precise mechanistic function of BRD8 remains somewhat unclear, especially in normal non-cancerous cells. At least, in embryogenesis, the accurate splicing of the *Brd8* transcript is required for correct post-germinal vesicle oocyte development ^51^. These studies nonetheless implicate BRD8 in developmental processes.

Several epigenetic pathways can be inhibited or promoted to increase the efficiency of the primed-to-naïve conversion. Inhibition of the H3K4 methylase KMT2A (MLL1) using a small molecule can promote the primed-to-naïve conversion ^52^. Indeed, we show that reduced *Brd8* expression led to reduced H3K4me1 at BRD8-bound loci, and also at the TSSs of primed-specific genes. Interestingly, BRD8 was bound to the promoter of *Kmt2a* in both ESCs and EpiSCs. However, the expression of *Kmt2a* was unaffected in the *Brd8* knockdowns and there was no overlap between genome-wide binding of BRD8 and KMT2A, suggesting that BRD8 and KMT2A affect the primed-to-naïve conversion independently. The transcriptional repressor *Sin3a* when overexpressed promotes the primed-to-naïve conversion ^53^, as does overexpression of the histone H3K27 demethylase *Kdm6b* (JMJD3) ^54^. Inhibition of the H3K79 methyltransferase DOT1L leads to a primed-to-naïve conversion with high efficiency ^20^. The TF *Zfp281* has been identified as a key primed transcriptional regulator that regulates *Tet1*, and thus DNA demethylation, to promote the primed state ^24, 55^.

ZFP281 also regulates EHMT1 and its catalytic target H3K9me to control ZIC2 genome-wide binding ^19^. There was a modest overlap of BRD8 binding with ZFP281 in EpiSCs, and this may partly explain the changes we see in H3K9me3 levels. Ultimately, these data all point to the idea that several epigenetic barriers are erected that impair the primed-to-naïve conversion. Enhancer loosening was seen previously in the primed to naïve-transition ^56^.

Potentially, the phenomenon we are seeing here is related to this process, as when *Brd8* was reduced we indeed saw a series of changes in chromatin accessibility at both primed and naïve genes, and a consequent change in enhancer chromatin marks (H3K27ac, H3K4me1). This was reflected in reduced KAT5 binding at primed-specific genes and at acetylated histones. These observations suggest enhancers are indeed weakened in response to reduced expression of BRD8.

Overall, our data supports a model for BRD8 in maintaining cell-type stability. BRD8 helps anchor the NuA4 complex to acetylated histones where it serves a role in maintaining active chromatin at promoters and transcribed gene bodies. Reduced *Brd8* expression disrupts KAT5 binding, leading to disrupted acetylation maintenance, and a consequent weakening of promoters and enhancers. This process occurs indiscriminately at both naive and primed-specific genes, but if the cells are exposed to a culture environment that favors conversion to a naïve state then the conversion is improved. Altogether, our data shows that BRD8 acts as a barrier for the conversion of the primed cells to the naïve state by helping to maintain a stable cell type through the maintenance of chromatin marks at key promoters.

## Acknowledgments

We thank Miguel A. Esteban (BGI, Shenzhen) for the gift of the OG2-MEFs and mice used in this study. We thank Xibin Lu for the professional support for FACS. We acknowledge the assistance of SUSTech Core Research Facilities. Funding was from the National Natural Science Foundation of China (32270597).

## Author contributions

L.S. designed the study, performed the majority of experiments, acquired funding, and helped prepare the manuscript, F.X. performed key experiments. G.M., Z.X. and Y.Z. performed experiments. L.S. performed some of the bioinformatic analysis. D.L and R.J contributed ideas and helped prepare the manuscript. A.P.H. designed the study, performed the bioinformatic analysis, acquired funding, supervised the study, and wrote the manuscript. All authors revised the manuscript.

