## Supplementary Information for "BRD8 guards the pluripotent state by sensing and maintaining histone acetylation"

#### Supplementary Tables

**Supplementary Table 1**. Table of naïve and primed-specific genes as defined as differentially expressed genes, with an absolute log2 (fold-change) of > 2.0 and a Bonferroni-Hochberg corrected p-value (q-value) < 0.01.

**Supplementary Table 2**. Summary of the Co-IP Mass spectrometry results

#### Methods

##### ESC and EpiSC cell culture and the conversion between the two cell types

EpiSCs were cultured feeder-free on dishes coated with fetal bovine serum (FBS; NATOCOR) in FA medium (N2B27 medium, 15 ng per ml bFGF (PeproTech) and 20 ng per ml activin A (PeproTech). N2B27-based medium: DMEM/F12 (HyClone) and Neurobasal (Gibco) mixed 1:1, supplemented with N2 (Gibco), B27 (Gibco), non-essential amino acids (Gibco), GlutaMAX (Gibco), sodium pyruvate (Cellgro), penicillin/streptomycin (HyClone), 0.1 mM β-mercaptoethanol (Gibco). Mouse EpiSCs were passaged using Accutase (Sigma) and seeded as single cells at approximately 50,000 cells in a well of a 6-well plate every 3 days. The medium was changed daily.

Mouse ESCs were maintained feeder-free on 0.1% gelatin in N2B27 medium supplemented with 2i/LIF at the following final concentrations: 1 μM PD0325901 (Sigma), 3 μM CHIR99021 (Sigma) and 1,000U per ml leukemia inhibitory factor (LIF; Millipore). Mouse ESCs were passaged by 0.05% trypsin-EDTA (Gibco) dissociation every 3 days.

To reprogram EpiSCs into rESCs, EpiSCs were dissociated into single cells using Accutase and plated at a density of 5,000 cells per well of 6-well plate coated with feeder in N2B27+FA medium supplemented with 5μM Y27632 (Selleck). The next day, N2B27+FA medium was changed into N2B27-2iL+Vc medium for 5 days, Vc (A4034, Sigma) was used at 50ug per ml. The medium was changed every day.

##### Cell transfection and shRNAs used in this study

Lentiviruses were generated from HEK293T cells using the jetPEI (Polyplus transfection). Mouse EpiSCs were infected with the individual lentiviral supernatant with the addition of Polybrene (Sigma) for 8 hours followed by selection with puromycin (Selleck) for 2 days before reprogramming.

shRNA inserts were cloned into pLKO lentiviral vectors. shRNA target sequences are listed as: shBrd8: 5'-GGTTCTTCCCATGATACATGG-3', shBrd1: 5’-CTAGAAGCTCAAGGGTATAAA-3’, shBrd2: 5’-TTATGTTCTCCAACTGCTAT-3’, shBrd3: 5’-GTATGCAGGACTTCAACACCAT-3’, shBrd9: 5’-TGGACCTGAGTTCACTGTCTA-3’, shBrdt: 5’-GCCAAGTCGACAAACAGCTATT-3’, shIgf2bp2-1:5’-CCGTTGTCAACGTCACCTATA-3’, shIgf2bp2-2: 5’-GCCGCATGATTCTTGAGATTA-3’,

**Co-immunoprecipitation and mass spectrometry**

Co-immunoprecipitation mass spectrometry (Co-IP/MS) was performed as previously described, (Zhang, et al., 2020). Briefly, proteins were extracted using NP-40 lysis buffer (50 mM Tris-HCl pH 7.5, 150 mM NaCl, 0.5% Nonidet P-40, 1 mM EDTA). Antibodies were pre-bound to 40 µl Dynabeads protein G or 40 µl protein A (10001D and 10003D, Life Technologies) for 4°C for 3-5 hours. Antibodies used include: 10 µg anti-BRD8 (A300-220A, Bethyl Laboratories). Dynabeads-antibody were mixed with proteins and rotated overnight at 4°C, then the protein-antibody-Dynabeads washed three times with lysis buffer. Proteins were detected on an Agilent 7700X.

Mass spec data was analyzed using MAXQUANT (Tyanova, et al., 2016). The resulting peptide matches were filtered using glbase3 (Hutchins, et al., 2014), using default filter settings four mouse proteins. Briefly, peptides detected by MAXQUANT were filtered based on several criteria. A protein was considered detected if it had at least 1 unique peptide and a minimum intensity of 1,000,000 (Razor+Unique). A protein was considered an interactor with BRD8 if the intensity was at least 2-fold above the anti-FLAG control. The resulting peptides were filtered to remove a list of common contaminating proteins (Briefly, protein names starting with: Rpl, Rps, Tub, Gapdh, Act, Myh, Ighg, Iglv, Col, Golga, Kif, Myl, Krt, Eif, Vim, Atp, Igkv, Ighv). The resulting table of filtered proteins is in **Supplementary Table 1**. In some figures the proteins were selected filtered based on their presence in the epigenetic factor or transcription factor databases EpiFactors and AnimalTFDB (Medvedeva, et al., 2015) (Hu, et al., 2019).

**Flow cytometry**

Cells were dissociated into single cells using Accutase and collected using centrifugation. After washing once with PBS, the cell pellet was resuspended with PBS containing 0.1% BSA, followed by filtration using a cell strainer (BD Biosciences) to remove large clumps of cells. The cells were then analyzed using an FACSCanto flow cytometer (BD Biosciences). The GFP fluorescence intensity was detected in the FITC channel. Data were analyzed with FlowJo (v10.8.1) software.

**Western blot**

After being dissociated and counted, 1×10$^6$ cells were collected and lysed in RIPA buffer (100µl) (Beyotime, P0013B) with protease inhibitor cocktail (Roche) on ice for 5min and boiled for 5min at 100 °C. The samples were separated using 10–12% SDS–PAGE and transferred onto a polyvinylidene difluoride membrane (Millipore) using a wet transfer system, and then incubated with the primary antibodies and secondary antibodies. The following primary antibodies were used: anti-H3 (Abcam, ab1791, 1:2000), anti-H3K27ac (Abcam, ab4729, 1:2000), anti-H3K9me3 (Abcam, ab8898, 1:2000), anti-H4acetyl (Millipore, 06-866, 1:2000); anti-BRD8 (Abcam, ab17969, 1:2000), anti-BRD8 (Bethyl, A300-220A, 1:2000), anti-β-actin (Sigma, A5541, 1:4,000), anti-KAT5 (Proteintech,10827-1-AP, 1:1000), anti-HA (haemagglutinin; Sigma H6908;1:1,000), anti-FLAG (Sigma, F1804, 1:5000).

**RNA-seq and analysis**

RNA-seq was performed as previously described (Hutchins, et al., 2017; Li, et al., 2017). Briefly, RNA was purified using RNAzol RT (RN190, MRC) according to the manufacturer's instructions. The samples were prepared for sequencing with RNA-seq NEB Next Ultra RNA Library Prep Kit (7530, NEB). Samples were sequenced on an Illumina Novaseq 6000.

RNA-seq data was aligned to the mouse genome using STAR (Dobin, et al., 2013), using the settings "--outFilterMultimapNmax 100 --winAnchorMultimapNmax 100 --outMultimapperOrder Random --runRNGseed 777 --outSAMmultNmax 1 --outSAMtype BAM Unsorted --twopassMode Basic --outFilterType BySJout --alignSJoverhangMin 8 --alignSJDBoverhangMin 1 --outFilterMismatchNmax 999 --alignIntronMin 20 --alignIntronMax 1000000 --alignMatesGapMax 1000000". Counts were assigned to features using scTE/te_counter (https://github.com/oaxiom/te_counter) against transcripts (GENCODE v32) or transposable elements (He, et al., 2021), and reads were GC normalized using EDASeq (Risso, et al., 2011). Differentially expressed genes were called using DESeq2 (Love, et al., 2014). Genes were considered differentially regulated if they had a q-value or 0.01 (Bonferroni-Hochberg corrected p-value) and a fold-change of 2 or 4 (specified in the appropriate figure legend). Gene ontology analysis was performed using GO-seq (Young, et al., 2010), and GSEA used fgsea (Sergushichev, 2016). Other analysis was performed using glbase3 (Hutchins, et al., 2014).

**ATAC-seq and CUT&Tag and their analysis**

ATAC-seq was performed as described (Buenrostro, et al., 2015; Li, et al., 2017). Briefly, nuclei from approximately 50,000 cells were extracted with lysis buffer (10 mM Tris-HCl pH 7.4, 10 mM

NaCl, 3 mM $MgCl_2$, and 0.2% (v/v) IGEPAL CA-630). Tagmentation reactions were performed *in situ* by addition of 50 µl reaction mix from a TruePrep DNA Library Prep Kit (TD502, Vazyme). DNA fragments were purified using the MinElute PCR Purification Kit (28004, Qiagen). ATAC-seq libraries were amplified with PCR for an appropriate number of cycles 18 and the sequence index was added by TruePrep® Index Kit V2 for Illumina (TD202, Vazyme). The amplified DNA libraries were purified using the VAHTS DNA Clean Beads (N411-02, Vazyme).

CUT&Tag was performed as described (Kaya-Okur, et al., 2019). Briefly, 1 × $10^5$ cells were washed twice with wash buffer (20 mM Tris-HCl pH 7.4, 150 mM NaCl, 0.5 mM spermidine and 1× protease inhibitors). Then cells were bound to concanavalin A beads (BP531, Bangs Laboratories) in binding buffer (20 mM Tris-HCl pH 7.4, 10 mM KCl, 1 mM $CaCl_2$ and 1mM $MnCl_2$). The bead-bound cells were washed once with buffer containing 0.01% digitonin, and primary antibodies were added to bead-bound cells in antibody buffer (4 mM EDTA and 0.2% BSA in wash buffer containing 0.01% digitonin) and rotated overnight at 4 °C. The primary antibodies below were used: anti-BRD8 (ab17969, Abcam), anti-KAT5 (10827-1-AP, Proteintech), anti-H3K27me3 (07-449, Millipore), anti-H3K4me1 (ab176877, Abcam), anti-H3K4me3 (ab8580, Abcam), anti-H3K27ac (ab4279, Abcam) and anti-H3K9me3 (ab8898, Abcam). Secondary antibody (ABIN101961, Antibodies-Online) was added to bead-cells-antibody in wash buffer and further incubated at RT for 1 h. pG-Tn5 (S603-02, Vazyme) was add to cells in '300-wash' buffer (20 mM Tris-HCl pH 7.4, 300 mM NaCl, 0.5 mM spermidine and 1× protease inhibitors, 0.01% digitonin) at RT for 1h. Tn5 was activated with 300-wash buffer supplemented with 10 mM $MgCl_2$ and washed five times with 500 µl 300-wash buffer. To stop the Tn5 reaction, 10 µl of 0.5 M EDTA, 3 µl of 10% SDS and 1.5 µl of proteinase K (20 mg/ml) were added 55 °C for 1 h. DNA fragments were extracted and PCR amplified using NEBNext HiFi 2× PCR Master mix (M0541L, NEB) using the settings: 72°C, 5 min, 98°C 30 s and 12-14 cycles of: 98°C 10 s, 63°C 10 s and 72°C 30 s. 300-500 bp DNA fragments were purified with the VAHTS DNA Clean Beads (N411-02, Vazyme). Libraries were sequenced by an Illumina sequencer.

For the ChIP-seq, ATAC-seq and CUT&Tag, these data contain a large number of Tn5 adaptors which were first trimmed using cutadapt. The data was aligned to the mouse mm10 genome using botwtie2 (Langmead and Salzberg, 2012), using the options '--very-sensitive --no-unal --no-mixed --no-discordant', and for ATAC-seq only the extra option '-X 2000' was also used. Aligned reads were filtered to include only correctly paired reads with a quality score > 20 (samtools view -F 1804 -q 20) that mapped to a standard chromosome. Peaks were detected using MACS2 (Zhang, et al., 2008), using the mouse genome and default parameters. Peaks analysis was unified using redefine_peaks, as described (Ma, et al., 2021). Epigenetic states were estimated using ChromHMM (Ernst and Kellis, 2017), using custom state models for

EpiSCs based on the epigenetic data generated in this study. Transcription factor motifs were detected using HOMER (Heinz, et al., 2010). Other analysis was performed using glbase3 (Hutchins, et al., 2014).

##### Data Availability and accession

Data generated in this study is available under accession number GSE253033. Several datasets were reanalyzed as part of this work. Including ChIP-seq data from the following studies: OCT4 ESC, SOX2 ESC, KLF4 ESC, TFCP2L1 ESC, ESRRB ESC (all GSE11431) (Chen, et al., 2008), PRDM15 ESC (GSE73692) (Mzoughi, et al., 2017), OTX2 EpiSC, ZIC2 EpiSC, SOX2 EpiSC, OCT4 EpiSC, OCT6 EpiSC (All GSE74636) (Matsuda, et al., 2017), MLL1 EpiSC (GSE73992) (Zhang, et al., 2016), and ZFP281 EpiSC (GSE93042) (Huang, et al., 2017). RNA-seq data from the following studies: PRJEB6168 (Yang, et al., 2017), GSE56096 (Buecker, et al., 2014), GSE58733 (Lowe, et al., 2015), GSE39656 (Klattenhoff, et al., 2013), PRJEB7132 (Takashima, et al., 2014), SRR1274703 (Factor, et al., 2014), GSE137627 (Huang, et al., 2020).

##### Statistics and Reproducibility

No statistical test was used to determine the sample size. The investigator was not blinded to the experimental details. Differential gene expression was calculated using DESeq2 (v1.36.0). A gene was considered significantly differentially regulated if it had an absolute fold-change of at least 2 and a Bonferroni-Hochberg corrected p-value (q-value) of <0.01. Gene ontology analysis was performed using goseq (v1.48.0) and statistics were calculated using goseq's internal statistical model. A gene ontology category was considered significantly enriched if there were at least 50 genes in that GO term and a Bonferroni-Hochberg corrected p-value (q-value) of <0.01. GSEA was performed using fgsea (v1.22.0). Gene sets were considered enriched or depleted if they had an absolute NES (normalized enrichment score) of at least 1.5 and a Bonferroni-Hochberg corrected p-value (q-value) of <0.01.

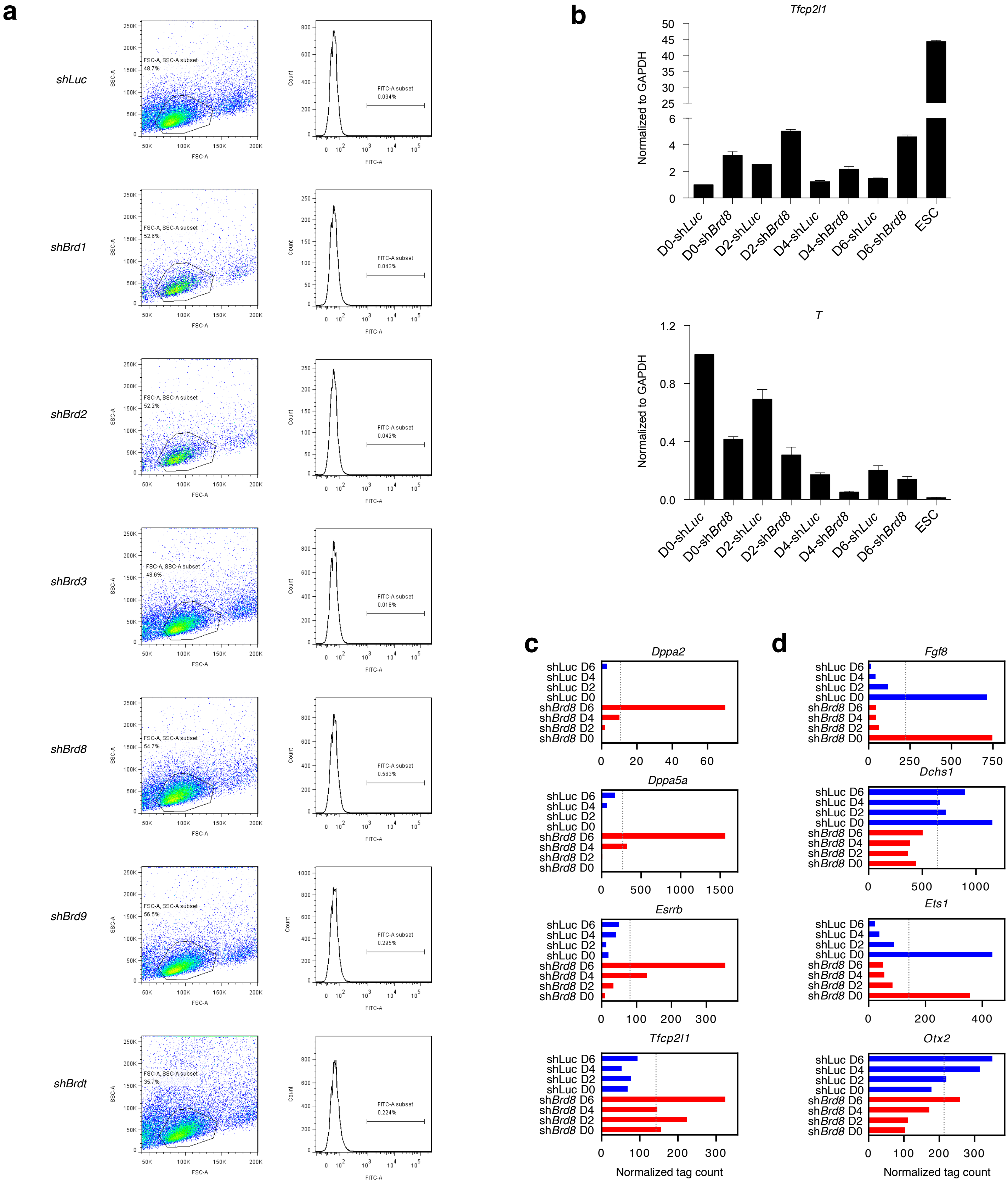


**Supplementary Figure 1**

**Supplementary Figure 1. BRD proteins impair the primed-to-naïve transition**

- **a.** Example flow cytometry histograms for EpiSCs at day 6 of the primed-to-naïve transition transfected with the indicated shRNAs. The left plots show the forward scatter (FSC) and the side scatter (SSC) and the gating strategy used. The right plots show the GFP+ histogram.
- **b.** Bar charts showing RT-qPCR for a naïve-specific gene (*Tfcp2l1*) and a primed-specific gene (*T*) in a primed-to-naïve transition cells transfected with a control shRNA against *Luc* or against *Brd8*. The experiment was repeated three times. Error bars are standard error of the means.
- **c.** Bar charts showing the expression levels of the indicated naïve-specific genes on days 0 to 6 of a primed-to-naïve transition. Cells were transfected with the indicated shRNA.
- **d.** Bar charts showing the expression levels of the indicated primed-specific genes on days 0 to 6 of a primed-to-naïve transition. Cells were transfected with the indicated shRNA.

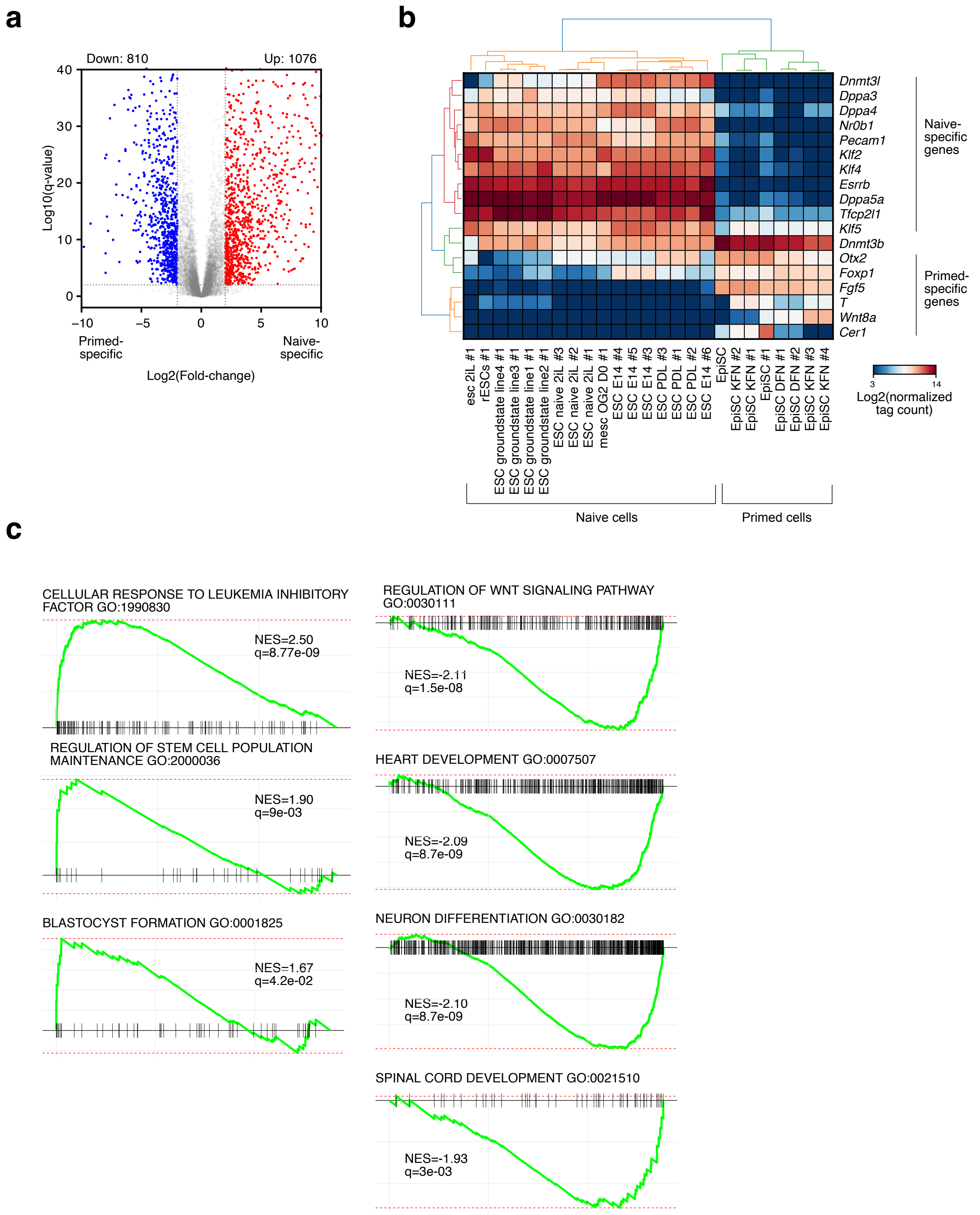


### Supplementary Figure 2

**Supplementary Figure 2. Reduced *Brd8* accelerates the primed-to-naïve transition.**

**a.** Volcano plot of a selection of EpiSC samples versus a selection of mESC samples. Genes were considered specific to naïve or primed if they had an absolute fold-change of at least 2.0 and a Bonferroni-Hochberg corrected p-value (q-value) of <0.01. RNA-seq data was reanalyzed from: PRJEB6168 (Yang, et al., 2017), GSE56096 (Buecker, et al., 2014), GSE58733 (Lowe, et al., 2015), GSE39656 (Klattenhoff, et al., 2013), PRJEB7132 (Takashima, et al., 2014), SRR1274703 (Factor, et al., 2014), GSE137627 (Huang, et al., 2020). Naïve cell types were defined as those grown in serum+LIF (SL) or in 2iL, whilst primed cells were defined as EpiSCs. See panel B for the specific samples.

**b.** Heatmap of a selection of naïve and primed-specific genes (from **panel A**), in the indicated RNA-seq samples.

**c.** Plots showing the enrichment of selected gene ontology terms for naïve-specific (left panels) or primed-specific (right panels) genes using GSEA. A term was considered significant if it had an absolute normalized enrichment score of at least 1.5 and a q-value of <0.01.

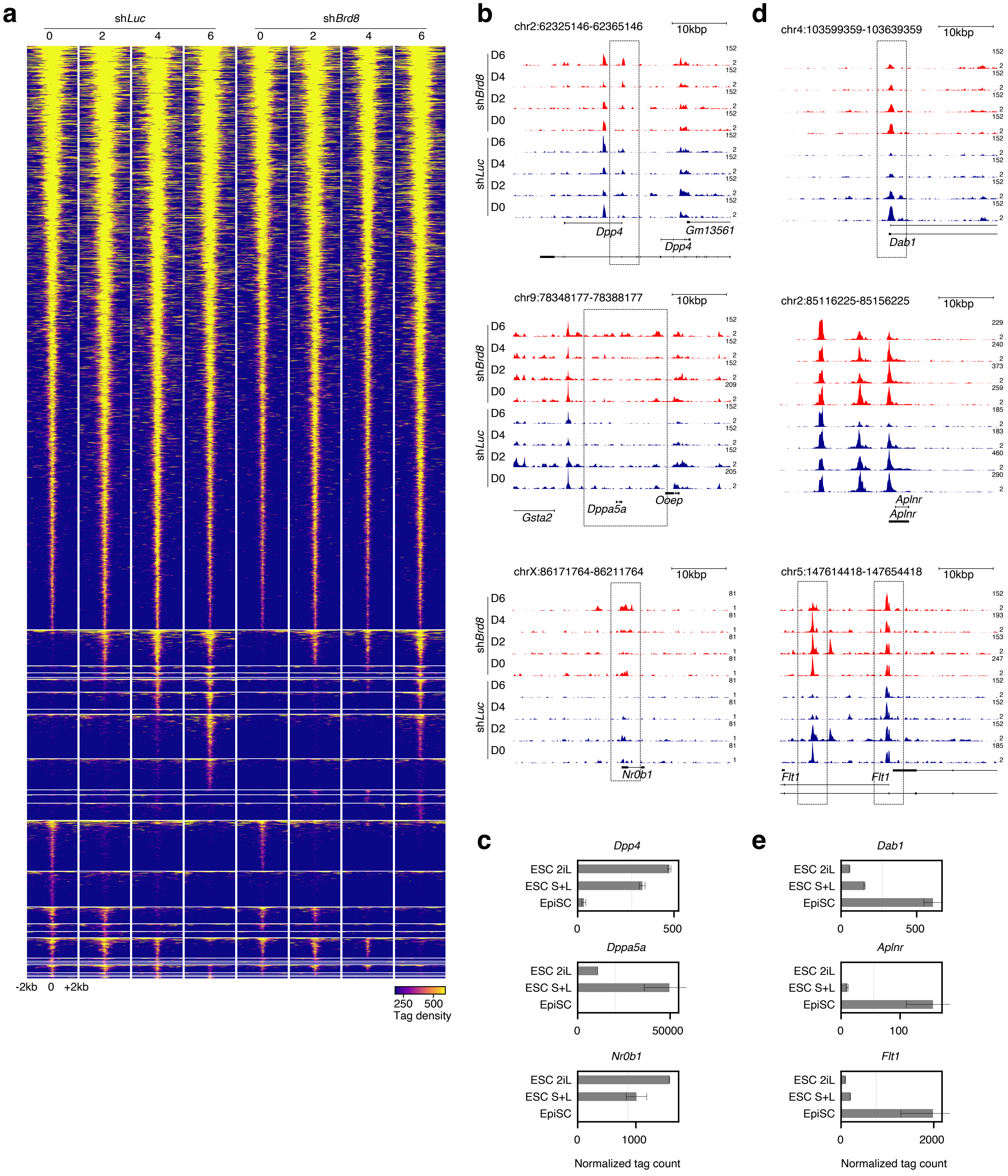


### Supplementary Figure 3

**Supplementary Figure 3. Chromatin accessibility during the primed-to-naïve transition.**

- **a.** Pileup heatmaps for ATAC-seq accessibility data showing the clusters of chromatin loci.
- **b.** Genome views at naïve-specific genes' loci showing the ATAC-seq data in EpiSCs undergoing a primed-to-naïve transition in cells transfected with the indicated shRNA.
- **c.** Bar charts showing gene expression or naïve-specific genes from RNA-seq data in ESCs ground in naïve conditions (2iL or SL) and primed (EpiSCs). Data is from **Supplementary Figure 2A**.
- **d.** Genome views at primed-specific genes' loci showing the ATAC-seq data in EpiSCs undergoing a primed-to-naïve transition in cells transfected with the indicated shRNA.
- **e.** Bar charts showing gene expression or naïve-specific genes from RNA-seq data in ESCs ground in naïve conditions (2iL or SL) and primed (EpiSCs). Data is from **Supplementary Figure 2A**.

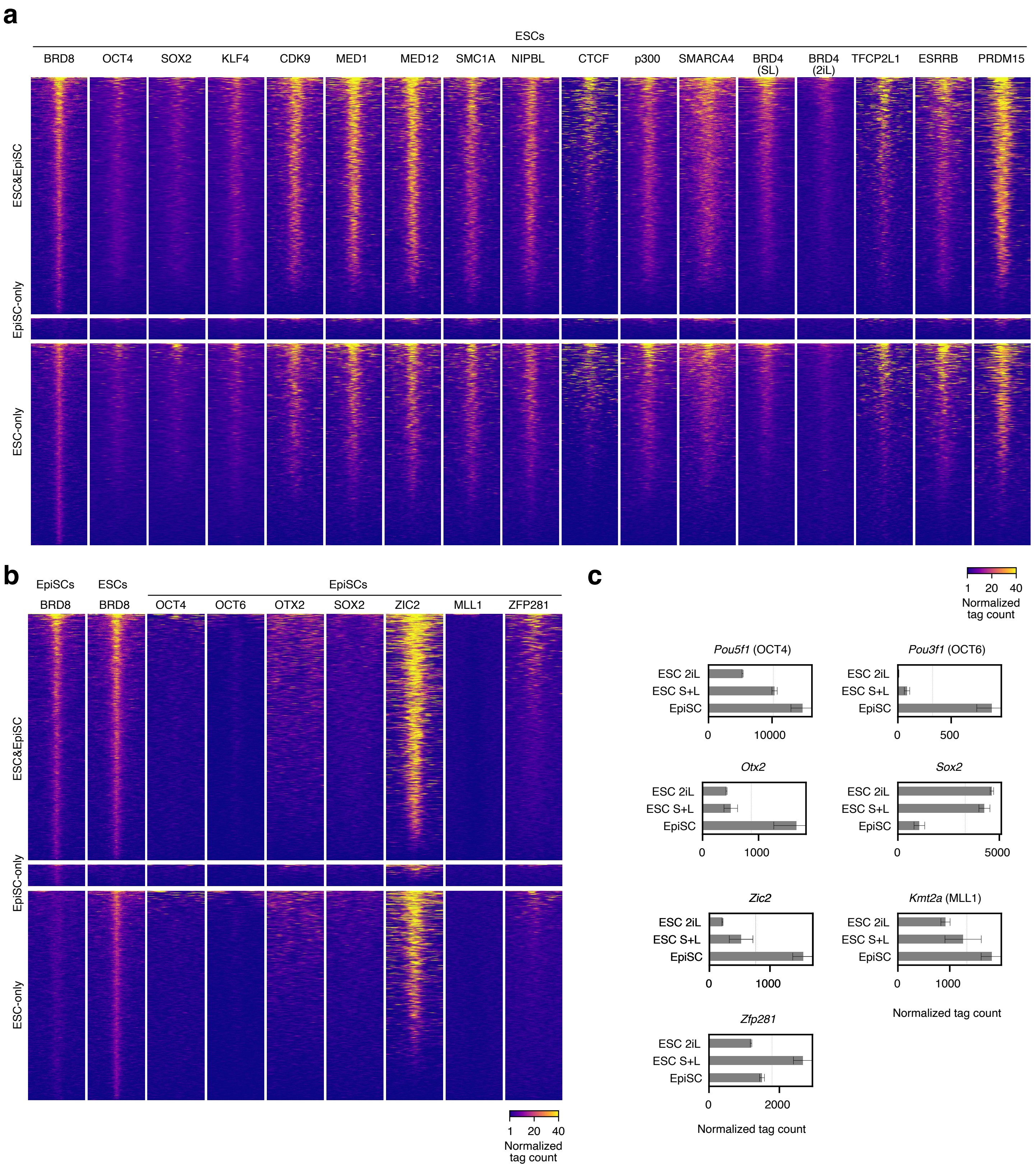


### Supplementary Figure 4

**Supplementary Figure 4. BRD8 co-binds with cell type-specific transcription factors in ESCs and EpiSCs.**

**a.** Heatmap pileups of binding at BRD8 loci in mESCs for selected transcription and epigenetic factors in mESCS. Data is from OCT4 ESC, SOX2 ESC, KLF4 ESC, TFCP2L1 ESC, ESRRB ESC (all GSE11431) (Chen, et al., 2008), PRDM15 ESC (GSE73692) (Mzoughi, et al., 2017).

**b.** Heatmap pileups of binding at BRD8 loci in EpiSCs (and BRD8 in ESCs). Data is from: OTX2 EpiSC, ZIC2 EpiSC, SOX2 EpiSC, OCT4 EpiSC, OCT6 EpiSC (All GSE74636) (Matsuda, et al., 2017), MLL1 EpiSC (GSE73992) (Zhang, et al., 2016), and ZFP281 EpiSC (GSE93042) (Huang, et al., 2017).

**c.** Bar charts showing gene expression of the transcription or epigenetic factors in **panel B** from RNA-seq data in ESCs ground in naïve conditions (2iL or SL) and primed (EpiSCs). Data is from **Supplementary Figure 2A**.

a

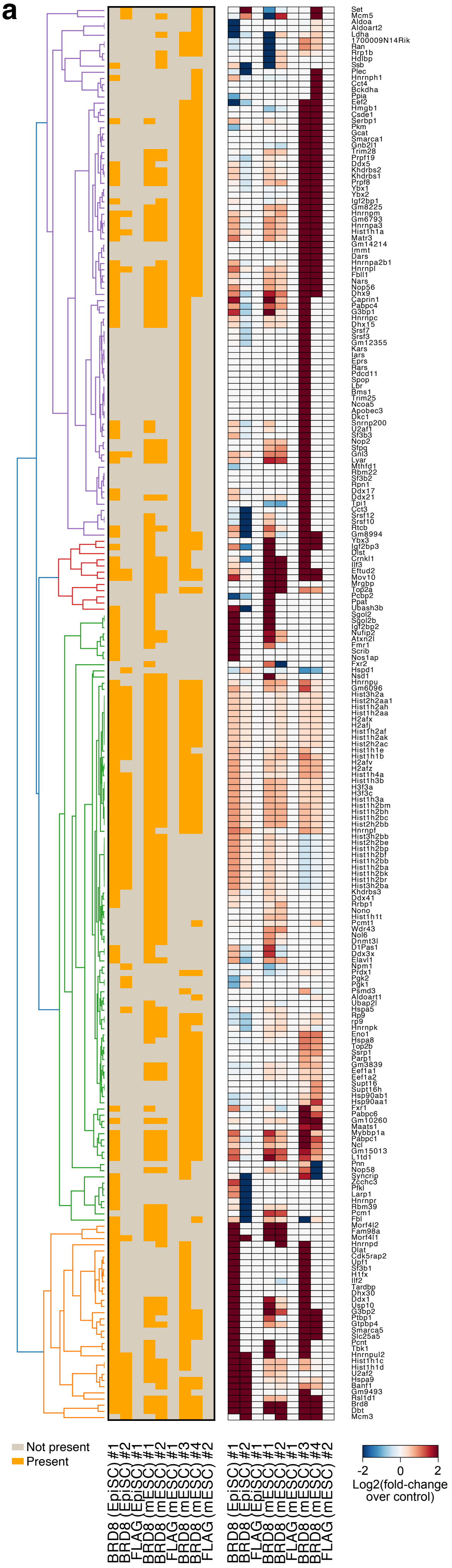


b

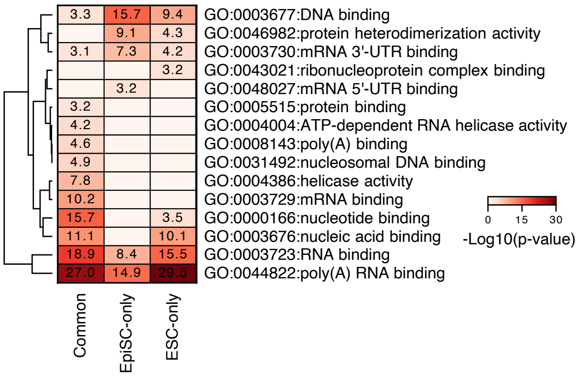


**Supplementary Figure 5**

**Supplementary Figure 5. BRD8 interacts with epigenetic factors in EpiSCs**

- **a.** Heatmap of the interacting proteins from Co-IP/MS data in EpiSCs and ESCs using antibodies against BRD8, or an anti-FLAG antibody as a control. The left heatmap shows the 'present' 'not present' call for the indicated protein in the indicated condition, whilst the right heatmap shows the detected protein's intensity. The full table is in **Supplementary Table 2**.
- **b.** Gene ontology (molecular function) for all proteins detected by Co-IP/MS (in specific cell types as defined in **Figure 4F**).

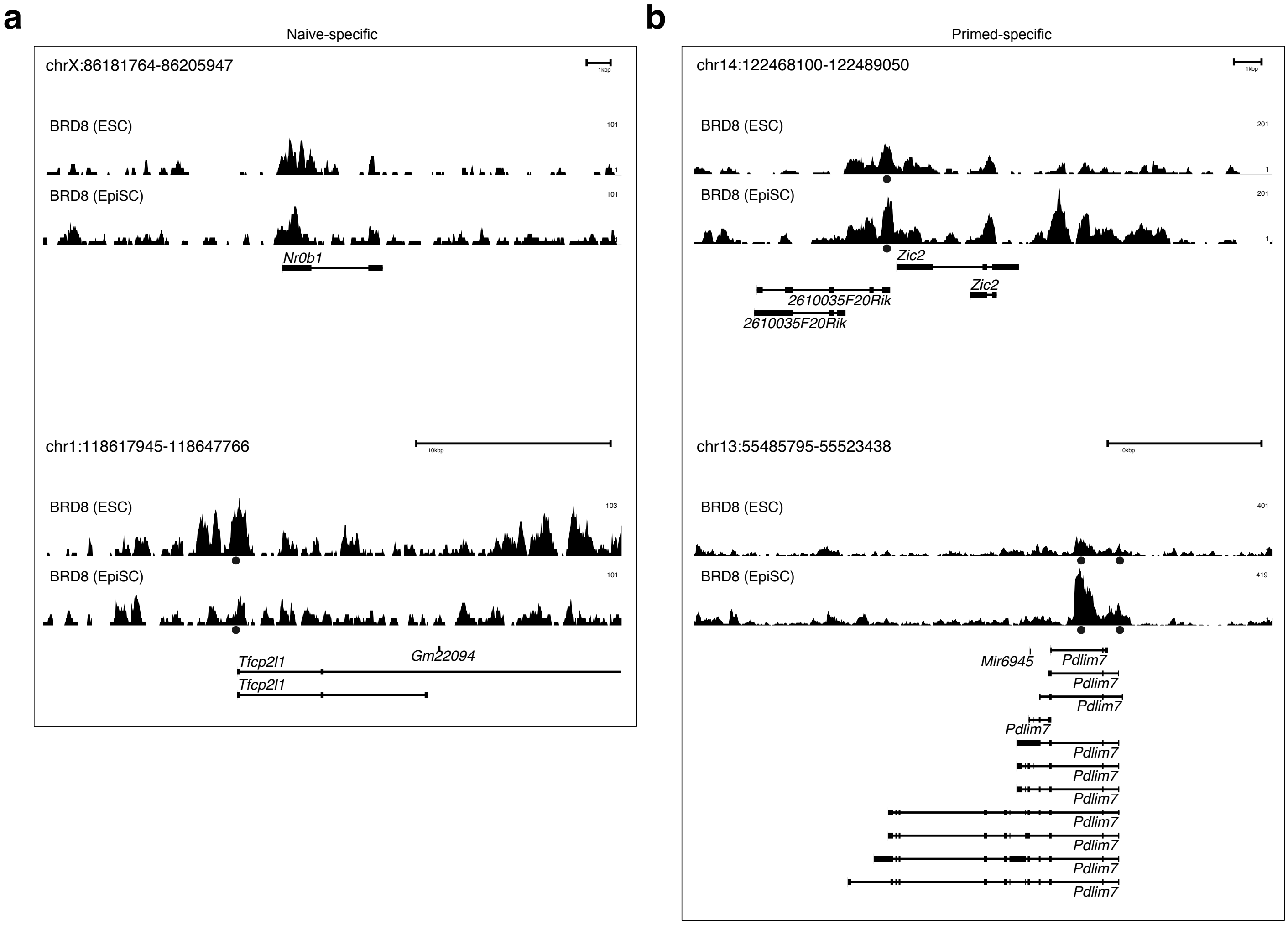


### Supplementary Figure 6

**Supplementary Figure 6. Example genome views at naïve and primed-specific genes.**

- **a.** Genome view of two naïve-specific genes, showing the BRD8 binding track (in black).
- **b.** Genome view of two primed-specific genes, showing the BRD8 binding track (in black).
